# Multipotent Progenitors Instruct Ontogeny of the Superior Colliculus

**DOI:** 10.1101/2023.04.16.537059

**Authors:** Giselle Cheung, Florian M. Pauler, Peter Koppensteiner, Thomas Krausgruber, Carmen Streicher, Martin Schrammel, Natalie Gutmann-Özgen, Alexis E. Ivec, Christoph Bock, Ryuichi Shigemoto, Simon Hippenmeyer

**Affiliations:** Institute of Science and Technology Austria (ISTA), Am Campus 1, 3400 Klosterneuburg, Austria; CeMM Research Center for Molecular Medicine, Austrian Academy of Sciences, Vienna, Austria; Medical University of Vienna, Institute of Artificial Intelligence, Center for Medical Data Science, Vienna, Austria; Department of Neurology and Neurological Sciences, Stanford University, Stanford, California USA

## Abstract

The superior colliculus (SC) in the mammalian midbrain is essential for multisensory integration, attention, and complex behavior (Basso and May, 2017; Cang et al., 2018). The mature SC cytoarchitecture is organized into distinct laminae and composed of a rich variety of neuronal and glial cell types (Ayupe et al., 2023; Edwards et al., 1986; May, 2006; Xie et al., 2021; Zeisel et al., 2018). Precise execution of the developmental programs regulating the generation of SC cell-type diversity is essential, because deficits due to genetic mutations have been associated with neurodevelopmental diseases and SC dysfunction (Jure, 2018; McFadyen et al., 2020). However, the fundamentals directing the ontogeny of the SC are not well understood. Here we pursued systematic lineage tracing at the single progenitor cell level in order to decipher the principles instructing the generation of cell-type diversity in the SC. We combined *in silico* lineage reconstruction with a novel genetic MADM (Mosaic Analysis with Double Markers)-CloneSeq approach. MADM-CloneSeq enables the unequivocal delineation of cell lineages *in situ*, and cell identity based on global transcriptome, of individual clonally-related cells. Our systematic reconstructions of cell lineages revealed that all neuronal cell types in SC emerge from local progenitors without any extrinsic source. Strikingly, individual SC progenitors are exceptionally multipotent with the capacity to produce all known excitatory and inhibitory neuron types of the prospective mature SC, with individual clonal units showing no pre-defined composition. At the molecular level we identified an essential role for PTEN signaling in establishing appropriate proportions of specific inhibitory and excitatory neuron types. Collectively, our findings demonstrate that individual multipotent progenitors generate the full spectrum of excitatory and inhibitory neuron types in the developing SC, providing a novel framework for the emergence of cell-type diversity and thus the ontogeny of the mammalian SC.

## MAIN TEXT

The mouse superior colliculus (SC) is located in the dorsal midbrain and essential for multisensory integration, attention, arousal brain states, and motor responses required for complex behavior (Basso et al., 2021; Cang et al., 2018; Oliveira and Yonehara, 2018). With six alternating strata of cell bodies and fibers, the superficial layers (sSC) are important for visual functions, receiving direct inputs from the retina; whereas the deep layers (dSC) are sites of auditory and somatosensory processing (Edwards et al., 1986; Ito and Feldheim, 2018; May, 2006; Sanes and Zipursky, 2020; Seabrook et al., 2017). SC dysfunction leads to deficits in sensory processing and has been implicated in neurodevelopmental diseases such as autism and attention deficit hypersensitive disorders (Jure, 2018; McFadyen et al., 2020).

The faithful production and distribution of the diverse neuronal and glial cell types during development is fundamental to the establishment of the highly complex SC cytoarchitecture, laminar arrangement and eventual circuit assembly. However, the cellular principles directing the generation of cell-type diversity and overall ontogeny of the SC remain poorly understood. While previous studies have identified radial glial progenitors (RGPs) lining the ventricular surface in the developing midbrain (Edwards et al., 1986; Gray and Sanes, 1991; Vanselow et al., 1989), their proliferation behavior and neurogenic potential is not known. Above all, the origins of SC excitatory glutamatergic and inhibitory GABAergic neuronal populations remain obscure despite that glutamatergic and GABAergic progenitor domains are thought to overlap in the developing dorsal midbrain (Achim et al., 2014; Arimura et al., 2019; Tan et al., 2002).

The SC is composed of a rich variety of distinct neuron types that show highly variable morphology, receptive field sizes, physiological properties, and synaptic target areas (Basso et al., 2021; Benavidez et al., 2021; Cang et al., 2018; Edwards et al., 1986). Yet, most information related to SC cell-type diversity has been derived based on physiological characteristics of a limited number of experimentally-accessible cells, primarily located in the superficial layers (Gale and Murphy, 2014; Masullo et al., 2019; Shang et al., 2015; Villalobos et al., 2018). Recent efforts have however systematically catalogued cell-types, based on single cell transcriptome, across the embryonic and adult midbrain (Ayupe et al., 2023; La Manno et al., 2021; Xie et al., 2021; Zeisel et al., 2018), and provide therefore an exciting starting point for our general understanding of the extent of SC cell-type diversity. Nonetheless, how cell-type diversity emerges from progenitor cells in the developing SC is not known. Here, we define the developmental principles governing RGP cell lineage progression, and the generation of cell-type diversity in the SC at single-progenitor cell resolution.

### A single common pool of progenitors in the SC

In order to obtain a global overview of cell-type diversity, and to temporally define *in silico* the emergence of neuronal cell types in the embryonic SC, we exploited a recent single-cell RNA-sequencing (scRNA-seq) dataset (La Manno et al., 2021) (Fig. 1a). We extracted a total of 26532 dorsal midbrain-specific single cells from embryonic (E) 9 to E18 time points (Fig. S1a). Uniform Manifold Approximation and Projection (UMAP), in combination with unsupervised clustering, identified a single continuum of cells including RGPs, immature and mature neurons (Fig. 1b and Fig. S1b). RGPs were highly abundant at E9 – E11 but sharply diminished by E12, coinciding with the peak of immature neuron abundance and the appearance of mature neurons (Fig. 1c). Within the neuronal population, we identified two largely-exclusive but continuous clusters of excitatory and inhibitory neurons, respectively, emerging simultaneously at E12 (Fig. 1d – 1e and Fig. S1b). The continuum of cells in the UMAP implied two distinct developmental trajectories from RGPs to neurons. Indeed, pseudotime analysis of the single-cell trajectory revealed connections from RGPs to mature neurons via immature neuronal states (Fig. 1f). The trajectory graph also identified a number of end points within the UMAP that correlated well with our unsupervised clustering analysis (Fig. S1c – S1d).

**Figure 1.**
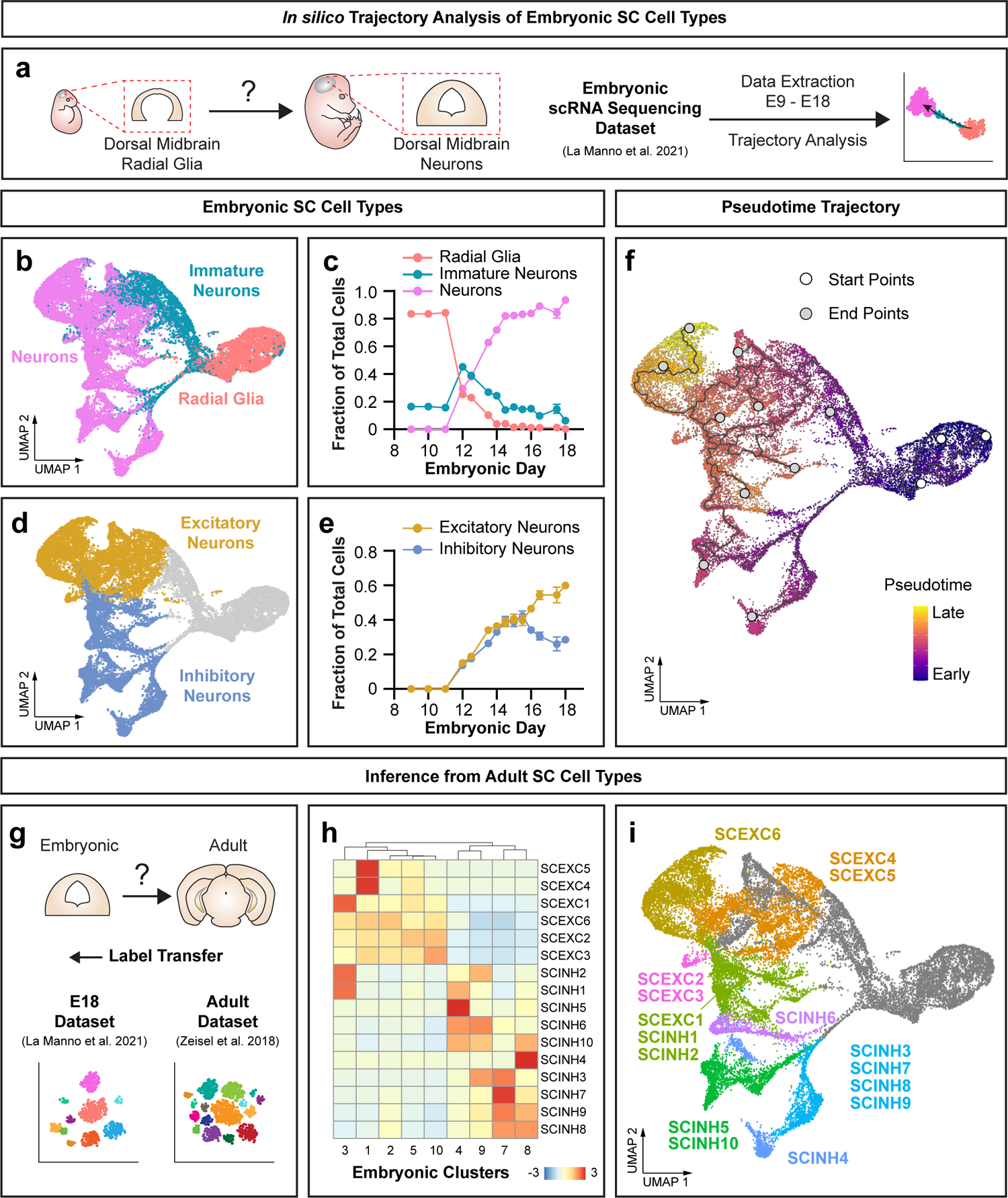
Emergence and developmental *in silico* trajectories of cell types in SC. **(a)** Schematic diagram illustrating the rationale and *in silico* analysis pipeline to assess the emergence of cell-type diversity in SC during embryogenesis, based on LaManno et al. dataset (La Manno et al., 2021). **(b-e)** UMAPs and line plots of relative cell abundance (fraction ± 95% Clopper-Pearson confidence intervals) for distinct clusters of radial glia, immature neurons and mature neurons (b and c); and excitatory and inhibitory neurons (d and e). **(f)** UMAP with coloration indicating pseudotime. Developmental trajectory is indicated with corresponding start (white) and end points (grey). **(g)** Schematic of label transfer analysis for inference from Zeisel adult dataset (Zeisel et al., 2018) to E18 dataset from La Manno. **(h)** Heatmap indicating the average similarity score of 9 embryonic neuronal clusters to 16 adult SC cell types extracted from Zeisel dataset. **(i)** UMAP illustrating the most similar adult SC cell types matching to individual embryonic clusters, coloring of cells according to Fig. S1c.

Next we assessed how the transcriptional profile of nascent SC neurons correlates with their mature state. We performed label transfer from an adult dorsal midbrain scRNA-seq dataset (Zeisel et al., 2018) to the E18 dataset (La Manno et al., 2021) (Fig. 1g). From the Zeisel dataset we extracted 16 neuronal cell types, comprising of 6 excitatory and 10 inhibitory neuron subtypes in the dorsal midbrain (Fig. S1e – S1f). In our analysis the individual mature SC cell types were matched to broader but specific clusters of embryonic neurons (Fig. 1h – 1i), indicating that a limited number of broad neuronal clusters in the embryo precipitates to all distinct mature cell types of the SC.

The above pseudotime trajectory analysis indicated a single population of RGPs from which all mature SC neuronal cell types seem to emerge. To approach such hypothesis we gained experimental access via genetic means, to the RGP population in the developing SC. We constructed a tamoxifen (TM)-inducible CreER driver line with transgene expression under the control of the *Frizzled-10* (*Fzd10*) promoter (*Fzd10-CreER^+/-^*) and confirmed targeted expression in embryonic dorsal midbrain progenitors (Fig. S2a – S2h). To more comprehensively assess the *Fzd10* cell lineage we crossed *Fzd10-CreER^+/-^* with the fluorescent *mTmG* reporter (Muzumdar et al., 2007) (Fig. S2i – S2k). We injected TM in *mTmG;Fzd10-CreER^+/-^*mice at E10.5, collected GFP^+^ cells of the *Fzd10* lineage at E12.5, E14.5, and E16.5 from the dorsal midbrain, and pooled samples for scRNA-seq using 10X Genomics technology (Fig. S2l). Our dataset including 5552 high-quality *Fzd10*-lineage single cells overlapped well with La Manno dataset in the integrated UMAP, and covered all midbrain-specific neuronal cell clusters in comparable proportions (Fig. S2m – S2r). Thus, our newly established *Fzd10-*CreER driver faithfully targets the relevant pools of embryonic RGPs giving rise to cell lineages comprising all the distinct mature neuronal cell-types in SC.

### Temporal lineage progression of SC progenitors

To precisely decipher ontogeny of SC cell-types from a single pool of RGPs we next pursued clonal analysis by labeling individual RGPs and following their lineages (Fig. 2a). We utilized Mosaic Analysis with Double Markers (MADM) technique to label individual units of clonally-related cells (Beattie et al., 2020; Contreras et al., 2021; Hippenmeyer et al., 2010; Zong et al., 2005). MADM relies on *Cre*-mediated interchromosomal recombination in dividing RGPs, thereby labeling the two daughter cells and their respective progenies in distinct red (tdT) or green (GFP) fluorescence colors (Fig. 2b and Fig. S3a). Given the quantitative nature based on single cell labeling *in situ*, MADM can provide a high resolution optical readout of progenitor proliferation behavior. To specifically target the SC, we used MADM reporter cassettes on chr.11 (Hippenmeyer et al., 2010) in combination with *Fzd10-*CreER driver.

**Figure 2.**
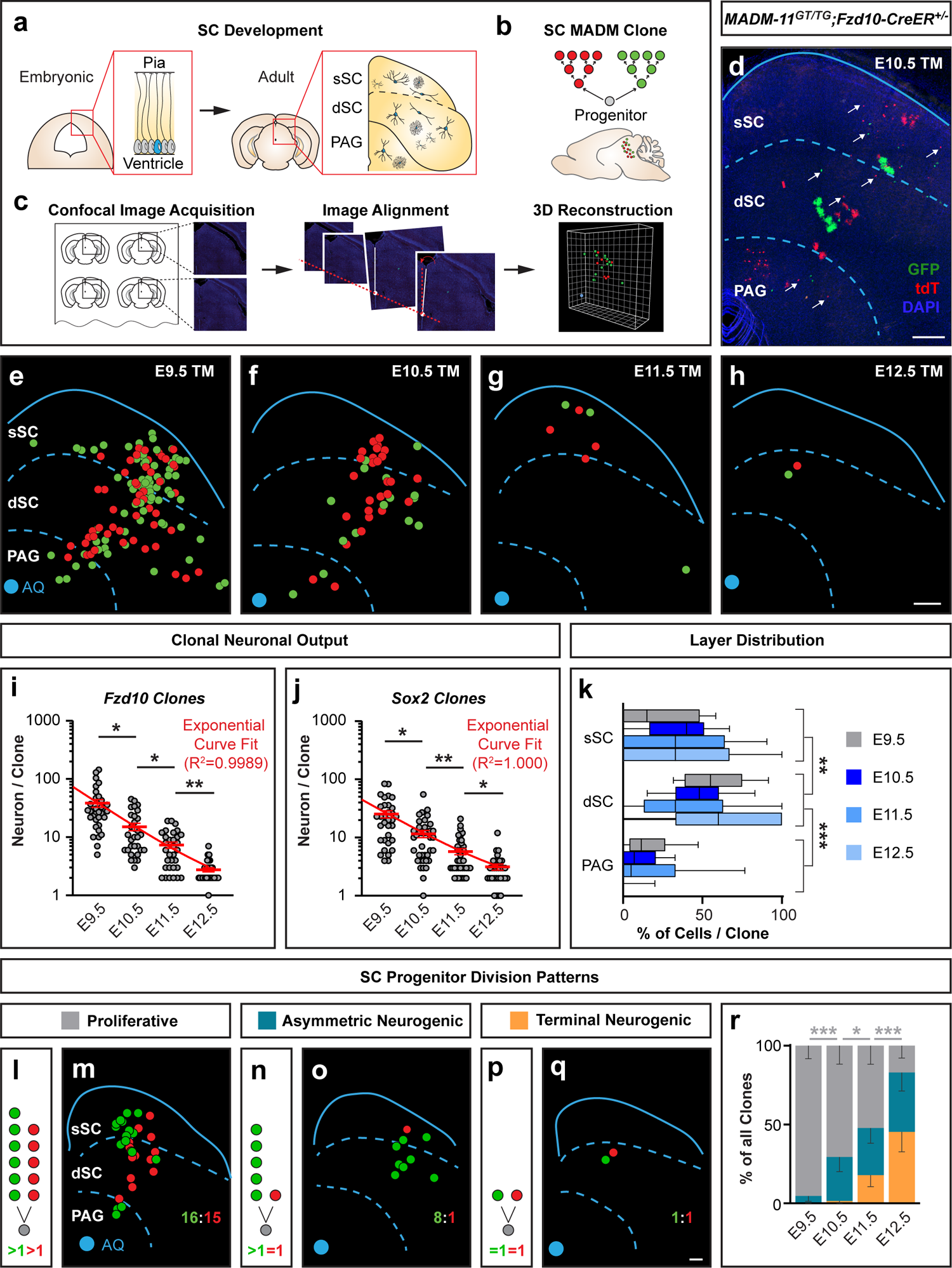
MADM clonal analysis reveals patterns of neurogenesis at single progenitor cell level in SC. **(a-c)** Schematic illustration of the emergence of cell-type diversity during development in SC (a); the appearance of differentially-labeled cell lineages in a MADM clone originating from an individual SC progenitor (b); and overview of analysis pipeline for image acquisition, alignment and 3D reconstruction of individual MADM clones (c). **(d)** Representative maximum z-projected image of a MADM clone in SC, induced at E10.5 and collected at P28 in *MADM-11^GT/TG^;Fzd10-CreER^+/-^*. Red and green neurons are marked by white arrows. Note that protoplasmic astrocytes and small oligodendrocyte clusters are also visible. **(e-h)** Representative reconstructions of MADM clones in SC, induced at E9.5 (e), E10.5 (f), E11.5 (g) or E12.5 (h), and collected at P28. Each dot represents the location of a single red or green neuron. The SC is outlined by a blue line and layers divided by blue dotted lines. Blue dot marks the location of the aqueduct (AQ). **(i-j)** Quantification of clonal neuronal output in *MADM-11^GT/TG^;Fzd10-CreER^+/-^* (*Fzd10* clones) (i), and in *MADM-11^GT/TG^;Sox2^CreER/+^* (*Sox2* clones) (j). The number of neurons per clone (mean ± SEM) across induction time points is plotted on the log_10_ scale. E9.5 (n=33), E10.5 (n=31), E11.5 (n=32), and E12.5 (n=30) in (i); E9.5 (n=33), E10.5 (n=39), E11.5 (n=39), and E12.5 (n=31) in (j). One-way ANOVA with Dunn’s post-hoc test; *p*=0.0113, *p*=0.0358 and *p*=0.0052 between consecutive time points in (i); *p*=0.0192, *p*=0.091 and *p*=0.0476 between consecutive time points in (j); *p*=0.7534, *p*>0.9999, *p*>0.9999, and *p*>0.9999 of the same time point at E9.5, E10.5, E11.5, and E12.5 between (i) and (j), respectively. Red line = exponential one phase decay curve fit of mean values. **(k)** Quantification of the percentage of neurons (box=median with 25-75 percentiles, whiskers=10-90 percentiles) located in each layer per clone across induction time points. Combined *Fzd10-* and *Sox2* clones at E9.5 (n=66), E10.5 (n=70), E11.5 (n=71), and E12.5 (n=61). Two-way ANOVA; *p*<0.0001 between sSC and dSC, *p*<0.0001 between dSC and PAG; *p*=0.9999, *p*=0.9999, and *p*=0.9999 between consecutive time points. **(l-q)** Illustration and a representative clone for each of three categories of SC progenitor division patterns: proliferative (l and m), asymmetric neurogenic (n and o), and terminal neurogenic (p and q). **(r)** Quantification of the proportion of all clones (mean + lower limits) in each category across induction time points; combined *Fzd10* and *Sox2* clones at E9.5 (n=64), E10.5 (n=68), E11.5 (n=67), and E12.5 (n=53). Fisher’s exact test comparing between proliferative and other clones; *p*=0.000164, *p*=0.034401, and *p*=0.0000642 between consecutive time points. \**p*<0.05; \*\**p*<0.01; \*\*\**p*<0.001. Scale bars = 200μm (d-h, m, o and q).

We induced MADM clones at E9.5, E10.5, E11.5, or E12.5 in *MADM-11^GT/TG^;Fzd10-CreER^+/-^*embryos and collected brains at postnatal day (P) 28-P30 (Fig. S3b) for analysis (Fig. 2c). We observed that SC MADM clones typically consisted of clusters of red and green neurons spanning across all layers of the adult SC, and the periaqueductal gray (PAG) (Fig. 2d and Fig. S3c – S3e). We analyzed 126 clones obtained from 131 *MADM-11^GT/TG^;Fzd10-CreER^+/-^* brains and observed that neuronal output of individual progenitors, measured by clone size, decreased exponentially over relatively short time window (Fig. 2e – 2i and Fig. S4a – S4d). Considerable clone size variability at all induction time points was observed. To independently validate and extend our dataset, we analyzed SC MADM clones (142 clones from 142 brains) that were induced with *Sox2-*CreER driver [uniformly expressed neural stem and progenitor cell (Arnold et al., 2011)]. We noticed comparable neuronal output patterns in *MADM-11^GT/TG^;Sox2^CreER/+^*brains corroborating our observations using *Fzd10-CreER^+/-^* (Fig. 2j and Fig. S4e – S4h).

SC MADM clones showed ‘cone-shaped’ architecture (Fig. S5a –S5i) and a disproportionate layer distribution of neurons (34 ± 2% in sSC, 51 ± 2% in dSC, and 15 ± 2% in PAG, Fig. 2k). The tangential dimension and overall dispersion of cells decreased significantly over time, resulting in narrower clones (Fig. S5j – S5l). In contrast, layer distribution of neurons (Fig. 2k) and the radial dimension of the clones (Fig. S5m) did not change over time. Thus, in contrast to other brain regions (e.g. neocortex), the emergence of laminae in the SC did not follow a temporally stereotyped pattern.

Next we determined the implicit cell division pattern of individual SC progenitors based on red and green MADM subclone size. We analyzed clones induced at progressively later time points from E9.5 to E12.5 and classified all clones into three categories: 1) Proliferative clones whereby the first division produced two self-renewing proliferative daughter cells; 2) Asymmetric neurogenic clones where the minority clone consisted of a single postmitotic neuron and the other subclone of more than one cell; and 3) Terminal neurogenic clones consisting of two single neurons labeled in distinct red/green colors (Fig. 2l – 2q). Based on quantitative assessment of the relative proportions of the three clone categories at each time point, RGPs quickly shift from proliferative to terminal neurogenic mode (Fig. 2r). Altogether, our MADM clonal analysis indicates that RGP lineage progression occurs rapidly in SC but that distinct laminae do not emerge in a temporally stereotyped pattern.

### MADM-CloneSeq reveals cell-type composition of individual clonal units

How do individual RGPs establish the full complement of neuronal cell diversity as observed in the mature SC? To address this issue we conceived a new approach, MADM-CloneSeq, combining the power of MADM in generating individual high confidence clonal units with scRNA-to determine the identity of clonally-related cells (Fig. 3a). Thus MADM-CloneSeq enables unprecedented correlation of lineage relationship and cell-type identity while preserving precise spatial information of SC neurons *in situ*.

**Figure 3.**
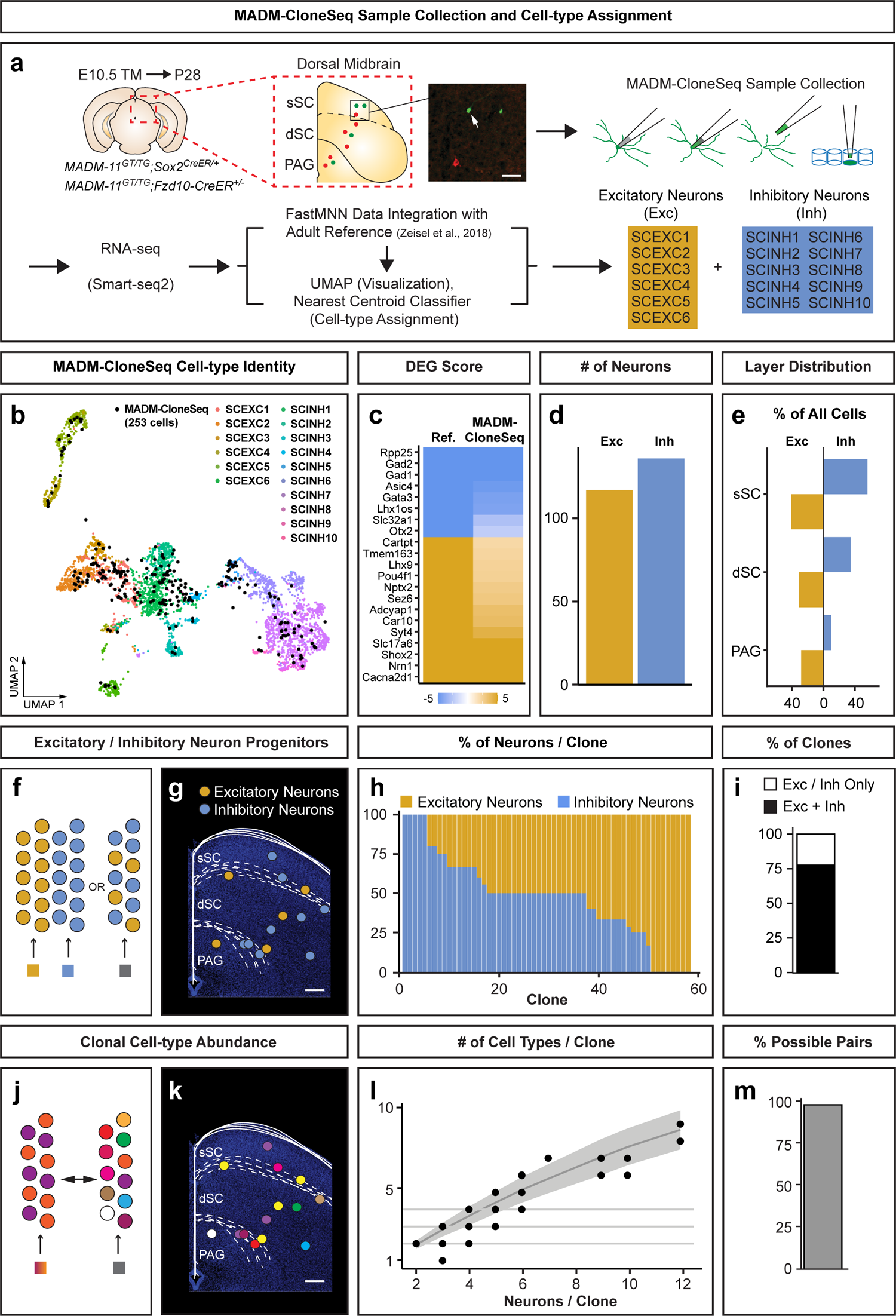
MADM-CloneSeq identifies multipotent progenitors with the capacity to generate all known neuronal types in SC. **(a)** Workflow of MADM-CloneSeq using acute brain slices containing MADM clones (either *Fzd10* or *Sox2* clones) in SC, induced at E10.5 and collected at P28. Each MADM-labelled cell was visualized by fluorescence and collected using a glass pipette and subjected to RNA-seq followed by data analysis. Informative cells were assigned to one of 6 excitatory or 10 inhibitory neuronal types according to reference Zeisel dataset (Zeisel et al., 2018). **(b)** UMAP indicating the overlay of 253 informative MADM-CloneSeq cells (black dots) onto reference neurons (colored dots). **(c)** Heatmap of DEG scores for selected genes expressed in excitatory- and inhibitory neuronal types in reference and MADM-CloneSeq datasets. **(d and e)** Quantification of the number of informative MADM-CloneSeq cells (d), and their relative layer distribution (e) for both excitatory-(n=117) and inhibitory (n=136) neuronal types. **(f-g)** Schematic diagrams illustrating two scenarios where individual SC progenitors either exclusively produce excitatory or inhibitory neuron types within a clonal unit, or have the capacity to generate both neuron types. The second scenario is illustrated with a hypothetical clone in (g). **(h)** Bar plot showing the relative proportion of excitatory and inhibitory neurons sampled in each individual MADM clone. **(i)** Quantification of the overall proportion of clones (n=58 combined *Fzd10* and *Sox2* clones) consisting of both, excitatory and inhibitory neuron types (black), or just one type (white). **(j and k)** Schematic diagrams illustrating two scenarios where individual SC progenitors generate either fixed or unrestricted clonal cell-type composition. The second scenario is illustrated with a hypothetical clone in (k). **(l)** The number of distinct cell types identified in each clone is plotted over the number of neurons sampled per clone (n=58 combined *Fzd10* and *Sox2* clones). Black line and grey ribbon indicate expected outcome for random cell type choice for each cell within a clone. Distribution of real data is not significantly different from randomized data at any clone size (z-score associated *p*-values=1 after multiple test correction). **(m)** The percentage of observed cell-type pairs in any given clone relative to the the total number of possible pairs considering 16 neuronal types (n=118 observed pairs out of 120 possibilities in combined *Fzd10* and *Sox2* clone dataset). Scale bars = 50μm (a), and 200μm (g and k).

By using glass pipettes we collected a total of 399 neurons from 87 SC MADM clones induced at E10.5, using either *Fzd10-* or *Sox2-*CreER driver, from acute brain slices at P28. After RNA-seq using the Smart-seq2 method, 253 informative cells from 58 clones passed quality filtering (Fig. S6a – S6g and Supplementary Table 1). We confirmed unbiased layer sampling (Chi-squared test; *p*=0.322; Fig. S6h). Our data analysis pipeline robustly assigned each MADM-CloneSeq cell to one of the 16 reference mature SC neuronal types (Zeisel et al., 2018), including 6 excitatory and 10 inhibitory neuron types (Fig. 3b and Fig. S1e – S1f). To verify precise cell-type assignment, we also showed that MADM-CloneSeq and reference cells shared highly similar marker gene expression when grouped into excitatory or inhibitory neuron types (Fig. 3c), or individual subtypes (Fig. S6i – S6j). Overall, we identified 117 excitatory and 136 inhibitory neurons in our dataset (Fig. 3d and Fig. S6k). While inhibitory neurons were most frequently sampled in the superficial layers (Chi-square goodness of fit: *p*=3e-4), excitatory neurons showed a more even distribution throughout the SC (Chi-square goodness of fit: *p*=0.2; Fig. 3e and Fig. S6l). Remarkably, all 16 neuronal cell types were detected among our MADM-CloneSeq cells with SCEXC2 being overall the most-, and SCINH4, 10 and 8 the least abundant cell types, respectively (Fig. S6m – S6n). Interestingly, we found that 13 out of 16 cell types were preferentially located in distinct layers with 7 in the sSC, 3 in the dSC and 3 in the PAG (*p*<0.1 z-score, Fig. S6o). Collectively, our MADM-CloneSeq data demonstrated that all known neuronal cell types in the SC are generated by resident RGPs.

### Individual progenitors in SC are multipotent

Next, we exploited the power of MADM-CloneSeq to delineate the lineage relationships between different SC cell types by analyzing the cell-type composition of individual SC clones. We obtained an average of 4 (ranging from 2 to 12) high-quality neurons in each clone (neurons/clone) which accounted for 60% (ranging from 31% to 100%) of all neurons/clone (Fig. S7a – S7b). To examine the lineage relationship between excitatory vs inhibitory neuron types, we first tested whether SC progenitors were restricted to producing one or the other type (Fig. 3f – 3g). Remarkably, we found that 78% of clones were composed of both excitatory and inhibitory neuron types. Thus we provide compelling evidence that glutamatergic excitatory and GABAergic inhibitory neurons originate from common progenitors in the developing SC (Fig. 3h – 3i and Fig. S7c). Clones consisting of only one type (i.e. excitatory or inhibitory) were significantly smaller (*p*=0.03, t-test) and included less informative MADM-CloneSeq cells (*p*=0.002, t-test). These results suggest that the probability of detecting both excitatory and inhibitory types was likely dependent on sample size (Fig. S7d – S7f). To determine the potential of SC progenitors beyond ‘simple’ excitatory and inhibitory property, we analyzed the abundance of all 16 neuron types within individual clonal units (Fig. 3j – 3k). We observed that the number of detected neuron types/clone increased with the number of informative cells/clone (Fig. 3l; black dots), with no significant difference to the randomized dataset (padj=1, z-score; Fig. 3l; grey line and shaded area). Furthermore, nearly all possible pairs of neuron types were present within individual clones (98% of 120 possibilities; Fig. 3m). Our findings thus suggest that individual RGPs have the potential to generate the complete spectrum of neuronal cell types in the developing SC. The information of the precise spatial location of sampled MADM-CloneSeq cells enabled us to assess lineage relationships between neuron types within and across SC layers. Using hierarchical clustering, we tested for any pattern of intra- or inter-laminar co-production of neuron types and found no significant preference (Fig. S7g – S7n). Taken together, we demonstrated that RGPs in SC are extraordinarily multipotent, with the capacity to produce the full spectrum of all, excitatory and inhibitory, neuron types without any pre-defined pattern related to location or their identity.

### *Pten* is cell-autonomously required for establishing cell-type diversity in SC

The multipotent nature of SC progenitors raises the key question of how defined relative proportions of different SC neuron types were established. To address this issue, we turned to investigate the role of candidate signaling pathways. Specifically, we focused on Phosphatase and Tensin homolog gene (*Pten*), since loss of *Pten* in other brain areas (i.e. neocortex) has been shown to affect the ratios of specific interneuron types (Vogt et al., 2015). However, a putative cell-autonomous *Pten* function in SC ontogeny has not been assessed at single cell level. We therefore utilized MADM technology and its exquisite single cell labeling property to probe the cell-autonomous function of *Pten* in controlling neuronal cell-type abundance in the SC. We generated genetic mosaics where *Pten* was sparsely deleted in RGPs, and compared the phenotype with wild-type RGPs in SC at P0 and P28 (Fig. 4a – 4c and Fig. S8a). To this end, we recombined a *Pten*-flox allele onto chr.19 containing the MADM reporter cassettes (see Methods) (Amberg and Hippenmeyer, 2021; Contreras et al., 2021). Next, we crossed *MADM-19^TG/TG,Pten-flox^* with *MADM-19^GT/GT^*;*Nestin-Cre*^+/-^ to generate *Pten*-MADM (*MADM-19^GT/TG,Pten^*;*Nestin-Cre*^+/-^) with *Pten*^-/-^ cells labeled with GFP and control cells with *Pten*^+/+^ tdT; and Control-MADM (*MADM-19^GT/TG^*;*Nestin-Cre*^+/-^; all cells *Pten*^+/+^) (Fig. S8b).

**Figure 4.**
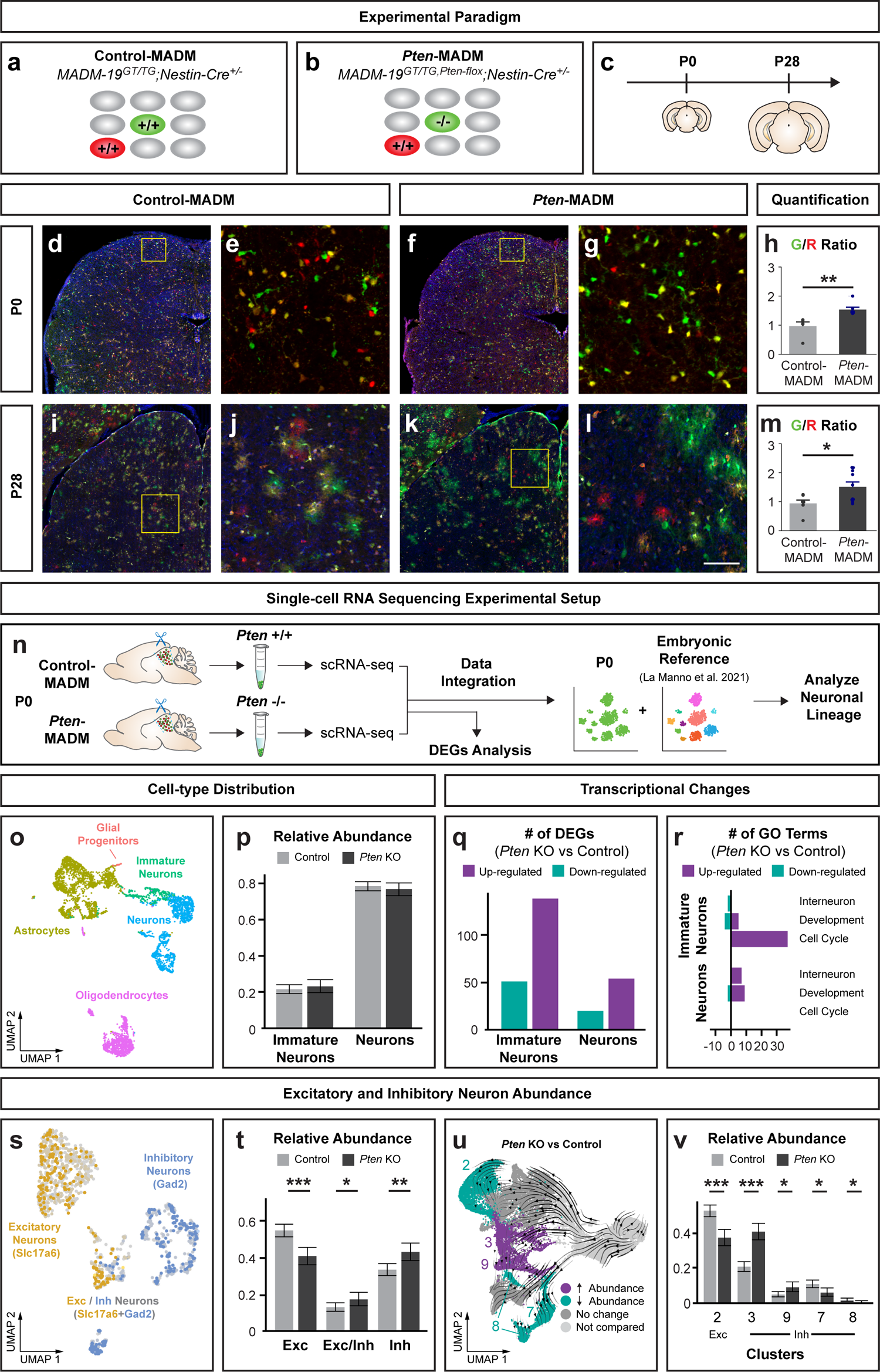
*Pten* is required for excitatory/inhibitory neuron cell-type balance in SC. **(a-c)** Experimental mosaic MADM paradigms to assess cell-autonomous *Pten* function in establishing cell-type diversity in SC. In Control-MADM (*MADM-19^GT/TG^;Nestin-Cre^+/-^*) (a), all cells were *Pten^+/+^* whereas in mosaic *Pten*-MADM (*MADM-19^GT/TG,Pten-flox^;Nestin-Cre^+/-^*) (b), red cells were *Pten^+/+^* and green cells *Pten^-/-^* according to the MADM principle. Sparse MADM labeling enables to probe the cell-autonomous function with single cell resolution in time course at P0 and P28 (c). See also Fig S8a-S8b. **(d-m)** Representative images (d-g and i-l), and quantification (h and m), of MADM-labelled cells in SC; at P0 (d-h), and P28 (i-m); in Control-MADM (d, e, i and j) and *Pten*-MADM (f, g, k and l). Magnified images of yellow boxed regions are shown in (e, g, j and l). Quantification of green/red ratio (mean ± SEM) of cells at P0 (h) and neurons at P28 (m) in Control-MADM: n=5 brains in (h) and n=7 sections in (m); *Pten*-MADM: n=7 brains in (h) and n=10 sections in (m). Mann-Whitney test; *p*=0.0025 in (h) and *p*=0.0431 in (m). \**p*<0.05; \*\**p*<0.01. **(n)** Experimental setup to isolate MADM-labelled green cells in Control-MADM (*Pten^+/+^*) and in *Pten*-MADM (*Pten^-/-^*) at P0 in dorsal midbrain for scRNA-seq and data analysis. Data integration was performed using La Manno embryonic reference dataset (La Manno et al., 2021). **(o)** UMAP indicating distinct clusters of neuronal and glial cell types in dorsal midbrain at P0 and identified based on La Manno annotations (n=2985 *Pten^+/+^* and n=1889 *Pten^-/-^* cells). **(p)** Relative abundance (fraction ± 95% Clopper-Pearson confidence intervals) of immature- and mature control (light grey, n=1080) and *Pten* KO (dark grey, n=567) neurons; chi squared *p*=0.49. **(q and r)** Number of up- or down-regulated DEGs (q) and their associated GO terms (r) in *Pten* KO compared to Control were plotted for immature and mature neuronal populations. GO terms for interneuron, developmental and cell cycle categories were indicated. **(s)** UMAP clusters containing either excitatory only, inhibitory only, or mixed neuronal populations of n=848 *Pten^+/+^* and n=436 *Pten^-/-^* cells. **(i)** 2. **(t)** Relative abundance (fraction ± 95% Clopper-Pearson confidence intervals) of each type was indicated between Control-(light grey) and *Pten* KO (dark grey). Chi squared statistics per category after multiple test correction; *p*=9.9×10^-06^ for Exc, *p*=0.048 for Exc/Inh, and *p*=0.0013 for Inh. **(u and v)** Cluster (as identified in Fig. S1c) assignment of Control- and *Pten* KO cells and relative abundance of Control-(light grey) and *Pten* KO (dark grey) cells in each cluster. Significant changes are highlighted on UMAP as higher/lower/unchanged in *Pten* KO relative to Control (u) and indicated as relative distribution (fraction ± 95% Clopper-Pearson confidence intervals) in bar plot (v). Chi squared statistics per cluster after multiple test correction; 9.5×10^-7^ for cluster 2, 4.4×10^-13^ for cluster 3, 0.014 for cluster 9, 0.014 for cluster 7, and 0.09 for cluster 8. \**p*<0.1 for; \*\**p*<0.01; \*\*\**p*<0.001. Scale bars = 200μm (d and f); 30μm (e and g); 400μm (i and k); and 90μm (j and l).

We first quantified the ratio of green to red cells in the SC at P0 (Fig. 4d – 4h) and P28 (Fig. 4i – 4m), and found a significant increase in *Pten*-MADM when compared to Control-MADM. Thus, the loss of *Pten* results in elevated overall SC neuron numbers, similar as observed in other brain areas (Groszer et al., 2001). However, whether the loss of *Pten* affects the numbers of specific SC neuron types is not clear. To resolve this issue we isolated green *Pten*^+/+^ and *Pten*^-/-^ cells from the dorsal midbrain at P0 in Control- and *Pten*-MADM mice, respectively, and subjected them to scRNA-seq using 10X Genomics technology (Fig. 4n).

After quality filtering and data integration with embryonic reference dataset (La Manno et al., 2021), we retained 2985 *Pten*^+/+^ and 1889 *Pten*^-/-^ cells with dorsal midbrain annotation (Fig. S8c – S8e). For downstream analyses, we focused on 1080 *Pten*^+/+^and 567 *Pten*^-/-^ neuronal cells. Our dataset showed that the relative distribution of immature and mature neurons was not altered by *Pten* deletion (Figure 4o - p). By comparing differential gene expression between *Pten*^+/+^ and *Pten*^-/-^ cells, we detected more differentially-expressed genes (DEGs) in immature than mature neurons (Fig. 4q). Gene Ontology (GO) enrichment analysis further revealed that *Pten*^-/-^ immature neurons showed upregulation of terms related to cell cycle (Fig. 4r and Fig. S8f), consistent with reported role of *Pten* in cell proliferation (Stiles et al., 2004). *Pten*^-/-^ mature neurons, on the other hand, were enriched in development and interneuron related GO terms (Fig. 4r and Fig. S8g). Our observation of an up-regulation of interneuron-specific genes suggested a potential alteration of cell-type composition upon deletion of *Pten*. To investigate this possibility, we focused on only mature neuronal populations (848 *Pten*^+/+^ and 436 *Pten*^-/-^ cells). UMAP and unsupervised clustering, together with marker expression analysis identified either excitatory only, inhibitory only or mixed neuronal clusters (Fig. 4s and Fig. S8h – S8j). We observed that the relative abundance of all three clusters was significantly altered upon *Pten* deletion, with a relative increase in inhibitory and decrease in excitatory neuronal clusters (Fig. 4t). To quantitatively assess whether all excitatory and inhibitory neuronal types were equally affected by *Pten* deletion we assigned cluster labels from our embryonic analysis (Fig. S1c) to each neuron. Next we determined the relative abundance of individual clusters between Control and *Pten* KO. We found that *Pten* deletion significantly altered the relative abundance of specific neuronal clusters (2, 3, 7, 8 and 9) while the rest (clusters 1, 4, 5, 10) remained unchanged (Fig. 4u - 4v). Among the affected clusters, the largest excitatory cluster (cluster 2) was decreased whereas the largest inhibitory cluster (cluster 3) was increased upon *Pten* KO. By superimposing velocities of developmental trajectories onto the UMAP of neuronal clusters (Fig. 4u), we revealed that the imbalance of excitatory and inhibitory abundance upon *Pten* deletion followed a trajectory-specific pattern. In the absence of *Pten* function, neuron clusters 3 and 9, which appear to originate from a distinct trajectory, became more abundant at the expense of neuron clusters 2, 7 and 8. Together, our results revealed an essential cell-autonomous *Pten* function in establishing appropriate quantitative balance of distinct excitatory and inhibitory neuron types, and thus overall cell-type diversity in SC.

## Discussion

Ever since the inaugural identification of RGPs in the mouse SC (Edwards et al., 1986) and optic tectum in non-mammalian vertebrates (Gray and Sanes, 1991; Vanselow et al., 1989) their neurogenic potential has not been determined. Whether resident RGPs constitute the sole source of SC neuron types is also not clear. In effect, in many brain areas the generation of excitatory and inhibitory neuron types is spatially segregated and nascent neurons often migrate long distances to reach their target area (Ayala et al., 2007; Evsyukova et al., 2013; Marin et al., 2010; Silva et al., 2019). For instance glutamatergic excitatory projection neurons in the developing neocortex emerge from RGPs located in the ventricular zone of the dorsal telencephalon (Kriegstein and Alvarez-Buylla, 2009; Lodato and Arlotta, 2015; Taverna et al., 2014) whereas GABAergic cortical inhibitory neurons originate from RGPs in the ventral ganglionic eminences (Bandler and Mayer, 2023; Lim et al., 2018; Wamsley and Fishell, 2017). Likewise, distinct neuron types destined for the olfactory bulb (Lim and Alvarez-Buylla, 2016; Lledo et al., 2008), ventral midbrain (Achim et al., 2012), or the cerebellum (Hatten, 1999) are generated at distant extrinsic progenitor niches. Here, our systematic cell lineage tracing approaches *in silico* and *in situ* at the single cell level revealed that all neuron types, including glutamatergic excitatory and GABAergic inhibitory neurons, in the developing SC emerge from local RGPs. At least at the qualitative level we demonstrate that no extrinsic progenitor source contributes to the diversity of cell-types in the SC.

The development of laminar brain structures including the cerebral cortex often follows temporally stereotyped patterns (Holguera and Desplan, 2018; Oberst et al., 2019). For instance, earlier-born corticofugal cortical projection neurons occupy lower layers and later-born callosal neurons locate to upper layers (Greig et al., 2013; Lodato and Arlotta, 2015). Interestingly, previous studies using 3^H^-thymidine autoradiography and analysis at the population level indicated possible inside-out order of SC lamina emergence (Altman and Bayer, 1981). However, our spatiotemporal analysis of individual clones with single cell resolution did not indicate any temporally stereotyped pattern of SC lamina appearance. Instead, SC neurons born at earlier stages did not exclusively populate the dSC and later born neurons were distributed across both, the sSC and dSC. Thus temporal RGP lineage progression did not predict the laminar position of SC neurons unlike in the developing cerebral cortex.

Our MADM-CloneSeq data comprising of precise quantitative and spatial cell lineage information, in combination with transcriptional identity as defined by scRNA-seq, demonstrates that all excitatory and inhibitory SC neuron types originate from common RGPs in a short temporal window. Importantly, individual RGP clones in mature SC did not show any pre-defined cell-type composition. In effect, the entire complement of all so far catalogued 16 distinct SC neuron types can emerge independently from single multipotent SC progenitors. These results suggest primarily intrinsic developmental programs of resident RGPs and provide definitive evidence of the principles instructing ontogeny and the emergence of cell-type diversity in SC.

Previous studies have revealed bi-potential progenitors located within the p2 domain in the developing spinal cord and generating both, excitatory v2aINs and inhibitory v2bINs (Peng et al., 2007) although these data emerged from population analysis rather than single clone assessment. Intriguingly, recent MADM-based clonal analysis revealed a population of embryonic *Sox2^+^* progenitors with the intrinsic potential to generate GABAergic Purkinje cells (PCs) and glutamatergic granule cells (GCs) in the cerebellum (Zhang et al., 2021). However, the generation of cerebellar inhibitory PCs and excitatory GCs is strictly separated in spatiotemporal dimensions. While PCs are generated around E12 by progenitors residing in the embryonic ventricular zone, GCs are produced only at postnatal stages when progenitor cells locate to the external granule cell layer (Butts et al., 2014). Thus the prolonged temporal progression of bi-potent cerebellar progenitor cells in combination with changing extrinsic signaling cues may significantly contribute to the sequential ‘reprograming’ of the cerebellar progenitor potential. Interestingly, in both spinal cord and cerebellar progenitor niches, Notch signaling appears to instruct the binary cell fate choice (Peng et al., 2007; Zhang et al., 2021). Yet, progenitor cells that can give rise to the full spectrum of neuron types, beyond binary excitatory/inhibitory neurotransmitter fate, have thus far not been identified in any brain area. Our findings therefore do not only provide a novel framework of SC ontogeny but extend our general understanding of progenitor cell lineage progression and potential.

Lastly, our analysis using MADM genetic mosaics identified a cell-autonomous role of *Pten* in SC neuronal cell-type abundance. Our findings suggest that while SC ontogeny stems from multipotent SC progenitors, the establishment of appropriate proportions of neuronal cell types is tightly regulated by *Pten*. Previous work has shown that deletion of *Pten* is associated with enlarged midbrain structures, which in turn disrupts sensory processing in mice (Clipperton-Allen et al., 2022; Ohtoshi, 2008). Intriguingly, conditional ablation of *Pten* in cortical interneurons results in a cell-type specific increase of parvalbumin/somatostatin (PV/SST) neuron ratio (Vogt et al., 2015). However, a general role for PTEN signaling in generating cell-type diversity has not been established. Based on sparse genetic mosaic analysis we show that *Pten* deletion results in an overall increase of *Pten^-/-^* mutant cells. Strikingly, the relative fractions of specific SC excitatory neuron types are significantly decreased whereas particular inhibitory neuron types are increased. The bias in SC neuron types emerged in a developmental trajectory-specific manner, implying critical *Pten* function at specific inferred cell lineage points during SC development. Altogether, our data demonstrate that *Pten* function is not only critical for balanced excitatory/inhibitory cell numbers but more generally for the faithful establishment of overall cell-type diversity in the developing SC.

## METHODS

### Mouse lines and maintenance

All animal procedures were approved by the Austrian Federal Ministry of Science and Research in accordance with the Austrian and European Union animal law (license number: BMWF-66.018/0007-II/3b/2012 and BMWFW-66.018/0006-WF/V/3b/2017). Experimental mice were bred and maintained according to regulations approved by institutional animal care and use committee, institutional ethics committee and the guidelines of the preclinical facility (PCF) at IST Austria. Mice were housed at 21±1°C ambient temperature and 40–55% humidity in 12 hrs dark/light cycles. Transgenic mouse lines with MADM cassettes inserted in Chr.11 (Hippenmeyer et al., 2010) and Chr.19 (Contreras et al., 2021), *Sox2^CreER^*(Arnold et al., 2011), *Nestin-Cre* (Petersen et al., 2002), *mTmG* reporter (Muzumdar et al., 2007), *Pten*-flox (Groszer et al., 2001) have been previously described.

*Fzd10-CreER^+/-^* transgenic mice were generated as follows. A 9.5 kb fragment containing upstream promoter region of mouse *Frizzled10* (*Fzd10*) gene was subcloned into pGEM vector (Promega). *CreER^T2^*gene (Feil et al., 1997), a gift from K. Miyamichi, was inserted in frame with the endogenous start ATG of *Fzd10*. Immediately downstream of *CreER^T2^*, a cassette comprised of an *IRES-taulacZ* followed by SV40 polyadenylation signal (Arber et al., 1999), gift from S. Arber, was inserted. The final *Fzd10-CreERT2-IRES-TLZ* transgene fragment was excised from the modified vector with *PmeI* and *AscI*, purified by agarose gel electrophoresis followed by QIAEX II (Qiagen), and injected into pronuclei of fertilized one-cell eggs from FVB at the Stanford Transgenic Facility. In total, seven *Fzd10-CreER^+/-^* transgenic founders were identified by PCR genotyping using *Cre*-specific primers. Upon assessment of TM (Sigma-Aldrich)-induced recombination in *Tau::mGFP* reporter (Hippenmeyer et al., 2005), one line (2#5) was maintained for all further investigation in this study.

All mouse lines were kept in mixed C57/Bl6, FVB and CD1 genetic background. In some experiments, wild-type CD1 mice were also used. Both male and female littermates of the desired genotypes were used randomly. All efforts were made to minimize the number of animals used following the 3R principles.

### Generation of *mTmG;Fzd10-CreER^+/-^*mice for scRNA-seq experiments

To selectively label and isolate cells of the *Fzd10*-lineage in the embryonic dorsal midbrain, *Fzd10-CreER^+/-^* mice were crossed with *mTmG* reporter line to generate *mTmG;Fzd10-CreER^+/-^* embryos. Timed pregnant females received intraperitoneal (IP) injections of TM (1 mg/mouse dissolved in corn oil) at E10.5. Upon *Cre* activation, a switch of fluorescent color from red (tdT) to green (GFP), according to the *mTmG* principle (Muzumdar et al., 2007) was observed in the embryonic midbrain cells (Fig. S2i – S2k). Dorsal midbrain from these *mTmG;Fzd10-CreER^+/-^* embryos were isolated at E12.5, E14.5 and E16.5 for scRNA-seq experiments.

### Generation of mice containing MADM clones in the superior colliculus

MADM clone induction in the SC was adapted from previously described protocols (Beattie et al., 2020; Hippenmeyer et al., 2010). In brief, *MADM-11^GT/TG^;CreER* mice were generated by crossing *MADM-11^GT/GT^;CreER* with *MADM-11^TG/TG^* mice (Fig. S3a). Two inducible *Cre* drivers (*Sox2*-CreER and *Fzd10-* CreER) were used independently to generate SC MADM clones.

To induce MADM clones, timed pregnant females were IP injected with TM (1-2 mg/mouse dissolved in corn oil) at E9.5, 10.5, 11.5, or 12.5, respectively. At E19, litters were delivered by caesarean section and raised with foster females. Experimental mice were collected for analysis between P28 – P30 (Fig. S3b). In total, we obtained 268 clones from 273 brains which equals to an overall average of 0.982 clone/brain (126 clones/131 *Fzd10-CreER^+/-^*brains and 142 clones/142 *Sox2^CreER/+^* brains).

### Generation of Control-MADM and *Pten*-MADM mice

To generate MADM genetic mosaic mice for *Pten* gene, we followed previously established protocols (Amberg and Hippenmeyer, 2021; Contreras et al., 2021). In brief, the *Pten-*flox allele was recombined onto Chr.19 containing the MADM-TG cassette to generate *MADM-19^TG/TG,Pten-flox^*stocks. Next, *MADM-19^TG/TG,Pten-flox^* were crossed with *MADM-19^GT/GT^*;*Nestin-Cre*^+/-^ to generate Control-MADM (*MADM-19^GT/TG^*;*Nestin-Cre*^+/-^) and *Pten*-MADM (*MADM-19^GT/TG,Pten-flox^*;*Nestin-Cre*^+/-^) mice (Fig. S8a). Upon *Cre* recombinase-mediated interchromosomal recombination for reconstitution of fluorescent MADM markers, all (red tdT^+^, green GFP^+^ and yellow tdT^+^/GFP^+^) cells in Control-MADM were wild-type (*Pten^+/+^*) whereas in *Pten*-MADM red tdT^+^ cells were wild-type (*Pten^+/+^*), green GFP^+^ cells homozygous mutant (*Pten^-/-^*) and yellow tdT^+^/GFP^+^ cells heterozygous (*Pten^+/-^*) in an otherwise unlabeled *Pten^+/-^* environment (Fig. 4a-c and Fig. S8b). Control-MADM and *Pten*-MADM brain samples were collected at P0 and P28 for histological studies; and at P0 for scRNA-seq experiments.

### Tissue collection for histological studies

Tissue collection for histological studies was performed according to previously described protocols (Beattie et al., 2020). For the collection of embryonic tissues, the embryos were dissected at specific developmental time points. P0 mice were collected on the day of natural birth. Brains were dissected after decapitation of mice and fixed in 4% PFA (paraformaldehyde; Sigma-Aldrich) overnight at 4°C. For the collection of postnatal tissues, mice at P28 – P30 were first anesthetized by a mixture of ketamine (65mg/kg), xylazine (13mg/kg) and acepromazine (2mg/kg) by IP injection. Transcardial perfusion of mice was performed with ice-cold PBS (phosphate-buffered saline) followed by ice-cold 4% PFA using a peristatic pump (Carl Roth, 4-6ml/min). Perfused brains were removed from the skull and post-fixed in 4% PFA overnight at 4°C. Upon complete fixation, both embryonic and postnatal brains were transferred to 30% sucrose solution (Sigma-Aldrich, dissolved in PBS) for 48-72 hrs. Tissues were embedded in Tissue-Tek O.C.T. (Sakura) and stored at −20°C or −80°C until further use.

For histological analysis, both embryonic and postnatal embedded tissues were cryosectioned using CryoStar NX70 cryostat (Thermo Fisher Scientific). Embryonic tissues were sectioned in either coronal or sagittal orientation at 20-30μm thickness and directly mounted onto superfrost glass slides (Thermo Fisher Scientific). Postnatal tissues were cryosectioned in coronal orientation at 30-45μm thickness and first collected in PBS and then mounted onto glass slides. For brains containing MADM clones in SC sections were collected, and kept in the same left-right orientation and in serial order, in multiple 24-well plates containing PBS. Mounted sections were air dried while protected from light and processed immediately for analysis.

### Antibodies and immunostaining of cryosections and acute brain slices

For immunostaining procedures, cryosections mounted on glass slides were first rehydrated with PBS at room temperature for 15 mins whereas acute brain slices (see below) were kept floating in PBS in multi-well plates to optimize permeability. Tissues were blocked for 2 hrs at room temperature in blocking solution. For cryosections we used 0.5% Triton X-100 with 5% Donkey Serum (Thermo Fisher Scientific) in PBS. For acute brain slices 1.2% Triton X-100 with 5% Donkey Serum in PBS was used. For some antibodies, an additional antigen retrieval incubation (Citrate Buffer; 192M citric acid + 0.05% Tween20; pH 6.0; at 85°C for 25 mins) was performed prior to adding blocking solution. After blocking, tissues were incubated for 16-48 hrs at 4°C with primary antibodies diluted in blocking solution. After washing with PBS with Triton X-100 (PBT), tissues were treated for 2 hrs at room temperature with secondary antibodies diluted in PBT. Finally, cell nuclei were stained with DAPI (4’,6-diamidino-2-phenylindole, Thermo Fisher Scientific, 1:5000 dilution) for 15 mins. Acute slices were mounted on glass slides and allowed to dry.

All sections were mounted using Mowiol 4-88 (Carl Roth) and 1,4-diazabicyclooctane (DABCO; Carl Roth) and stored at 4°C until image acquisition. The following primary antibodies were used: chicken anti-GFP (Aves, GFP1020, 1:500), rabbit anti-RFP (MBL, PM005, 1:500), goat anti-tdTomato (SICgen, ab8181-200, 1:500), chicken anti-beta galactosidase (Abcam, ab9361, 1:1000), rabbit anti-Ki67 (Abcam, ab15580, 1:500), rabbit anti-DCX (Abcam, ab18323, 1:200), and rabbit anti-olig2 (Millipore, AB9610, 1:200). The following secondary antibodies were used: donkey anti-chicken-Alexa488 (Jackson Immuno, 705545447, 1:1000), donkey anti-rabbit-Alexa568 (Life Technologies, A10042, 1:1000), donkey anti-goat-Alexa568 (Invitrogen, A11057, 1:1000), donkey anti-rabbit-Alexa647 (Life Technologies, A31573, 1:1000) and donkey anti-goat-Alexa647 (Life Technologies, A21447, 1:1000).

### Image acquisition and processing

Before image acquisition of sparsely-labelled MADM clones in SC, mounted serial sections were first screened for clones using an axioscope (Zeiss Axio Imager, Zeiss) coupled to a CoolLED p300 SB light source (CoolLED) and equipped with Plan-Apochromat 10x/0.45 and 20x/0.8 objectives (Zeiss). Green and red fluorescence were observed using an HC-dualband GFP/DsRed filter (F56-420, AHF). The presence of MADM-labelled cells in the SC was documented for subsequent confocal image acquisition. Confocal image acquisition was performed using LSM 800 series inverted confocal microscopes (Zeiss) and analyzed using ZEN Blue 2.3 and 2.6 software (Zeiss). Confocal images were acquired in z-stacks and tiles with excitation lasers 405, 488, 561, and 640nm. Plan-Apochromat 10x/0.45 and 20x/0.8 objectives were used. In order to allow accurate post-acquisition image alignment of clones spanning across multiple serial sections, the entire dorsal midbrain hemisphere consisting of the SC, PAG, midline and aqueduct were included in the image. Tiled images were subsequently stitched and exported in .tif format as either z-stacks or orthogonal projections using ZEN Blue built-in processing functions.

### MADM Clone 3D reconstruction

Serial confocal z-stacks of each MADM clone in SC were first aligned in Fiji software v1.53 (Schindelin et al., 2012) using custom-generated macro *3DSerialSectionAlignement.ijm*. The macro relied on user input to define a line ROI in the middle of each stack to be aligned against the previous stack. User input included the manual drawing of a straight line from the dorsal most border of the aqueduct along the midline using the *Straight line tool*. Upon execution of the macro, the first stack was rotated such that the midline was exactly vertical. Subsequent stacks were rotated in the same manner and then shifted in the x- and y-dimension such that the aqueduct aligned with the previous stack. The macro automatically saved individual aligned stacks, an aligned stack of the complete clone and its maximum z-projection. After image alignment, initial data analysis involved counting of the number of red and green cells per clone. Next, the 3D coordinates of each cell were marked using ImageJ *Point tool* and saved in *ROI manager*. For each clone, a reference point (x=0, y=0, z=0) was always set to the dorsal most point of the aqueduct in the middle z-plane of the clone. Apart from the 3D coordinates of each cell, other parameters were also noted: color (belonging to the red or green subclone), morphology (neuron, astrocyte, or oligodendrocyte), spatial location (sSC, dSC and PAG visibly demarcated by DAPI signal). For population analysis, astrocytes were identified based on their distinct protoplasmic morphology and oligodendrocytes were labelled using anti-Olig2 antibodies to distinguish them from small neurons. All information were then stored in Microsoft Excel for further analysis. For the quantification of 3D parameters of clones we determined a midline projecting through the reference point and the centroid of the clone. The maximum radial displacements of cells from the reference point in 3D were then calculated for each clone. Similarly, maximum tangential displacement of cells from the midline in 3D was also determined for each clone. The cell dispersion in each clone was calculated based on the standard deviation of each cell location in 3D. For MADM population analysis of Control-MADM and *Pten*-MADM tissues, cell counting was performed using a built-in plugin *Cell Counter*. The counts and x-y coordinates of red and green cells were exported to Microsoft Excel for further analysis.

### Preparation of single cell suspension for scRNA-seq

Embryonic dorsal midbrains were dissected at E12.5, E14.5 and E16.5 by first cutting the skin and skull to expose the dorsal part of the brain. An incision was made between the forebrain-midbrain boundary and another one between the midbrain-hindbrain boundary with fine surgical scissors. Two horizontal cuts were then made through the aqueduct to free the dorsal midbrains. In total, 9, 8, and 3 dorsal midbrains were pooled for each of E12.5, E14.5 and E16.5 time points, respectively.

To isolate P0 dorsal midbrains, mice were first decapitated and the brains dissected from the skull. After removing the cortices to expose the midbrain, two cuts were made with sharp razor blades: one between the forebrain-SC boundary and another between the SC-inferior colliculus boundary. Both cuts were angled to meet at the aqueduct thus avoiding contamination from ventral midbrain regions. In total, 26 Control-MADM and 24 *Pten*-MADM dorsal midbrains were separately pooled. The subsequent preparation procedures of single cell suspension from these tissues were adapted and modified from previous protocols (Laukoter et al., 2020). Dissected tissues were first incubated in Earle’s Balanced Salt Solution (EBSS, Thermo Fischer Scientific) containing Papain (Eubio) and DNaseI (Eubio) for 30 mins at 37°C with gentle shaking at 150rpm. After adding Ovomucoid I-Albumin (Eubio), tissue suspension was briefly dissociated mechanically with a pipette. After centrifugation at 1000rpm for 10 mins at room temperature, cell pellet was re-suspended and further mechanically dissociated until a homogeneous cell suspension was obtained. A second centrifugation at 1500rpm for 10 mins at room temperature produced the final pellet for fluorescence-activated cell sorting (FACS) for embryonic samples. P0 samples were processed further to remove debris due to an increased amount of myelination in the midbrain at this time point. Cell pellets were re-suspended and mixed with cold Debris Removal Solution (Miltenyi Biotec). The mixture was carefully overlaid with cold PBS before centrifugation at 3000 x g for 10 mins at 4°C. The top two layers containing PBS and debris were then aspirated and removed. The remaining cleaned cell suspension was then collected by a last centrifugation at 1000 x g for 10 mins at 4°C for FACS. Cell sorting for all GFP-labelled single cells was performed immediately using a SH800 Cell Sorter (Sony). In total, 33,000 labelled cells were collected from each embryonic time point while 14,000 and 12,000 labelled cells were sorted from P0 time point of Control-MADM and *Pten*-MADM, respectively. All cells were sorted in freshly prepared cold DMEM (Thermo Fisher Scientific) supplemented with fetal bovine and horse serum (Thermo Fisher Scientific) and processed immediately for scRNA-seq.

### Acute brain slice preparation and screening for MADM clones for MADM-CloneSeq

*MADM-11^GT/TG^;CreER* experimental mice containing MADM clones in SC at P28–P30 were first deeply anesthetized via IP injection of a mixture of ketamine (90mg/kg) and xylazine (4.5mg/kg), followed by transcardial perfusion with ice-cold, oxygenated (95% O_2_, 5% CO_2_) artificial cerebrospinal fluid (ACSF) containing (in mM): 118 NaCl, 2.5 KCl, 1.25 NaH_2_PO_4_, 1.5 MgSO_4_, 1 CaCl_2_, 10 Glucose, 3 Myo-inositol, 30 Sucrose, 30 NaHCO_3_ prepared in diethyl pyrocarbonate (DEPC; Sigma-Aldrich)-treated water; pH=7.4. The brain was rapidly dissected and coronal slices of 200 µm thickness including the SC were cut using a Linear-Pro7 vibratome (Dosaka, Japan). To preserve left-right orientation, each brain was marked on the ventral part of one hemisphere with a clean razorblade. Individual slices containing SC were first screened on both sides for the presence of MADM-labelled cells using an inverted fluorescence phase-contrast microscope (BZ-9000E; Keyence, Japan). Images were captured and processed using BZ-II Viewer (Keyence, Japan) for documentation and also used post-hoc to locate cells for collection. Screened slices containing SC MADM clones were returned to a custom-made multi-well recovery chamber (designed and 3D-printed by Robert Beattie, https://www.printables.com, 3D model 361319) and left to recover for 20 mins at 35°C, followed by a slow cool down to room temperature over 40–60 mins.

### MADM-CloneSeq sample collection

The sample collection steps for MADM-CloneSeq were adapted from previously published protocols (Cadwell et al., 2020; Cadwell et al., 2017) and optimized for speed and coverage of sparsely labelled cells of MADM clones. To prevent RNA degradation, all solutions were prepared using water pre-treated with DEPC. All work surface and equipment were cleaned using RNase away solution (Thermo Fisher Scientific) prior to experiments. Single acute slice containing MADM clones was first transferred to a BX51WI microscope (Olympus, Tokyo, Japan) and superfused with ACSF at a rate of 1–2 ml/min at room temperature. Individual neurons were visualized under a 60x objective via infrared / differential interference contrast (IR/DIC) video system using an XM10 camera (Olympus) and cellSens software (Olympus). To observe red or green fluorescence, light was emitted from a pE-300 LED light source (CoolLED, Andover, UK). Glass pipettes (B150-86-10, Sutter Instrument, Novato, CA) previously treated with DEPC and dried at 80°C were pulled using a P-1000 pipette puller (Sutter Instrument) to generate pipettes with opening of around 3-5μm or 1-3MΩ. Immediately before use, each pipette was filled with 3μl of pipette solution which consisted of 5% RNase inhibitor (Takara) in RNase-free PBS (Invitrogen) filtered through a 0.22μm filter. Red or green fluorescent neurons were approached with the pipette under positive pressure to avoid tissue contamination of the pipette tip. Each cell was approached from around 45° angle above the surface. When contact with cell membrane was made, tight seal formation was achieved under IR/DIC visualization. The complete aspiration of the cell body was immediately performed by applying a gentle and steady suction while monitoring under fluorescence and typically takes 5–10 s. The negative pressure was removed immediately once the target cell was collected and the pipette was carefully withdrawn from the acute slice to avoid contamination. The content of each pipette was mixed with 1μl of 0.8% Triton-X (final concentration of 0.2% Triton-X, Sigma-Aldrich) and then stored in individual wells of a 96-well plate which was kept at −80°C until cDNA library preparation. At the end of sample collection, all acute slices which contained the whole or part of a clone were fixed in 4% PFA at 4°C overnight. Fixed acute slices were then washed in and transferred to PBS and stored at 4°C until batch immunostaining procedures to reconstruct clones.

### MADM-CloneSeq and single-cell RNA-sequencing

cDNA libraries were prepared from single MADM-CloneSeq cells, followed by Smart-seq2 protocol (Picelli et al., 2014) using custom reagents (VBCF GmbH). Pools of libraries were sequenced on Illumina platforms at the VBCF NGS Unit (https://www.viennabiocenter.org/facilities/).

Control-MADM, *Pten*-MADM, and pooled embryonic dorsal midbrain samples containing *Fzd10*-lineage cells were processed for scRNA-seq. cDNA libraries were generated using the Chromium Controller and the Next GEM Single Cell 3’ Reagent Kit (v3.1, 10x Genomics) according to the manufacturer’s instructions. Single cell suspensions were incubated with Zombie NIR fixable viability dye (Biolegend) and viable GFP^+^ cells were sort-purified on a Sony SH800 cell sorter. The samples were processed individually according to the manufacturer’s protocol. Libraries were sequenced by the Biomedical Sequencing Facility at the CeMM Research Center for Molecular Medicine of the Austrian Academy of Sciences, using the Illumina NovaSeq 6000 platform. Raw sequencing data was pre-processed and demultiplexed using bcl2fastq (v 2.20.0.422).

### Embryonic scRNA-sequencing data analysis

Initial analysis, including integration was performed using R v4.1.2 and Seurat v4.1.0.

#### La Manno et al. dataset

For the embryonic reference single cell count data (La Manno et al., 2021) was downloaded from https://storage.googleapis.com/linnarsson-lab-loom/dev_all.loom. Loom files were processed using loomR (0.2.1). Cells were retained based on the following annotations stored as column attributes: Tissue: Midbrain or MidbrainDorsal, Subclass contains: midbrain or mixed. For cells with Age e9.0, e10.0, e11.0 we retained cells with the following Location attributes: Caudal and dorsal midbrain (caudal m1A), Alar plate of the midbrain and diencephalon, Roof plate of midbrain, diencephalon and pallium, Midbrain-hindbrain boundary diffuse, Midbrain and diencephalon roof plate, Midbrain alar plate, Midbrain roof plate, subpallium ganglionic eminence, Midbrain diencephalon roof plate, Midbrain dorsal, hindbrain lateral, Midbrain. Class: Neuron, Neuroblast, Radial glia. A Seurat object was prepared with these cells using CreateSeuratObject with min.cells = 5, min.features = 500 parameters. We split this data by the Age attribute, normalised each dataset using NormalizeData with normalization.method = “LogNormalize”, scale.factor = 10000 parameters and determined variable features using FindVariableFeatures with selection.method = “vst”, nfeatures = 2000 parameters.

#### Pooled embryonic Fzd10-lineage data

Initial analyses was performed using cellranger-7.0.0 with transcriptome version: mm10-2020-A, including intronic information. All downstream analyses were performed using filtered_feature_bc_matrix.h5 file from CellRanger. We created a Seurat object from this data using CreateSeuratObject with min.cells = 3, min.features = 200 parameters and retained only high quality cells based on these parameters: nFeature_RNA > 200 & nFeature_RNA < 6000 & percent.mt < 5. Normalisation and finding variable features was done as for the reference.

All reference and the pooled embryonic *Fzd10*-lineage data were integrated using SelectIntegrationFeatures, FindIntegrationAnchors and IntegrateData with default parameters. For downstream analyses we used the predictive power of the large, integrated dataset, if not stated otherwise. Visualizations focused on La Manno data only, to improve clarity of data presentation. All subsequent analyses were performed with R v4.2.1 and Seurat v4.2.0.

#### UMAP, cluster assignment

Initial clustering was performed using ScaleData, RunPCA (npcs = 30), RunUMAP (reduction = “pca”, dims = 1:26), FindNeighbors (reduction = “pca”, dims = 1:23), FindClusters (resolution = 1.8). We identified 2 clusters with low numbers of detected genes that were also disconnected from other cells on the UMAP. Cells from these 2 clusters were removed and the data re-clustered: RunPCA (npcs = 50, verbose = FALSE), RunUMAP(reduction = “pca”, dims = 1:30), FindNeighbors (reduction = “pca”, dims = 1:30), FindClusters(resolution = 0.2). Final clustering is visualised in Fig. S1c.

#### Developmental trajectory

Fig. S1b: Gene expression was visualized using FeaturePlot with order = T, min.cutoff = “q25”, slot = “scale.data” parameters. Fig. 1b, d, f, i: UMAPs show only La Manno et al. cells. Fig. 1b: Colors were assigned based on the Class attribute. Note that Neuroblasts are labeled as Immature Neurons. Fig. 1c:

Relative abundances of La Manno et al. cells with a Class attribute were calculated at each available developmental stage. Note that Neuroblast was renamed to Immature Neurons. Fig. 1d: Seurat clusters as shown in Fig. S1b were assigned a cell type using cells from LaManno et al. and the corresponding Class and the Subclass attributes. Clusters assigned as Excitatory Neurons or Inhibitory Neurons were colored. Fig. 1e: Relative abundances of La Manno et al. cells with a Subclass attribute containing glutamatergic (Excitatory Neurons) or GABA (Inhibitory Neurons) were calculated at each available developmental stage. Error bars for both figures were determined with clopper.pearson.ci (alpha = 0.05, CI = “two.sided”) from GenBinomApps v1.2. Fig. 1f, Fig. S1d: For monocle3 (v1.2.9) analysis the Seurat object was converted using as.cell_data_set (SeuratWrappers v0.3.1), cells were clustered using cluster_cells and the trajectory graph was learned using learn_graph with learn_graph_control = list(ncenter=1000, nn.k=25, geodesic_distance_ratio=0.25, euclidean_distance_ratio=1.5), close_loop = T, use_partition = T parameters. Indicated start points of trajectories were defined manually using order_cells with reduction_method = “UMAP” parameter.

#### Inference of adult cell types

Fig. S1f, Fig. 1h - 1i: We extracted UMI counts of relevant cells from the adult mouse superior colliculus (Zeisel et al., 2018) and prepared a Seurat object as described for MADM-CloneSeq reference cells. Using this Seurat object we followed the standard workflow: NormalizeData, FindVariableFeatures (selection.method = “vst”, nfeatures = 2000), ScaleData, RunPCA (npcs = 30), RunUMAP (reduction = “pca”, dims = 1:20). The resulting UMAP is shown in Fig. S1f. We reasoned that the latest embryonic time point (e18) will be most similar to adult cell types. Therefore we extracted raw UMI counts for e18.0 cells from the embryonic reference and performed the following processing workflow: CreateSeuratObject (min.cells = 3, min.features = 200), NormalizeData, FindVariableFeatures(selection.method = “vst”, nfeatures = 2000). For label transfer, first transfer anchors were determined: FindTransferAnchors with dims = 1:20, reference.reduction = “pca”. With these anchors label transfer was performed: TransferData (dims = 1:20). This analysis defines similarity scores for each e18 embryonic reference cell to each of the adult cell clusters. Note that we focused this analysis on neurons and thus excluded clusters 0, 6, 11 (Fig. S1c) from this analysis. For Fig. 1h we calculated the mean similarity score of embryonic reference cells in each Seurat cluster (Fig. S1c) to each of the adult cell types. Heatmap of the resulting matrix was drawn using pheatmap (v1.0.12) with scale=“row”. Fig. 1i: Finally we inferred for each adult cell type the most likely cluster of origin in the embryonic data by the highest mean similarity score and visualised this association on the UMAP.

#### Fzd10-transgene characterization

Fig. S2m-S2o: For QC plots we used nFeature_RNA as genes, nCount_RNA as transcripts. Fig. S2q, S2r: Number of cells from embryonic reference or from pooled embryonic *Fzd10*-lineage cells was counted in each Seurat cluster as shown in Fig. S2q (NOTE: only *Fzd10*-lineage cells are shown in this UMAP) and Fig. S1c. Relative number of cells in each cluster were calculated and plotted. Pearson’s product-moment correlation and p-value were calculated using cor.test function.

### MADM-CloneSeq data analysis

#### Reference dataset

As a reference dataset we used published single cell RNA-seq data (Zeisel et al., 2018) by downloading the raw read counts in loom format (https://storage.googleapis.com/linnarsson-lab-loom/l5_all.loom). All statistical analyses were performed in R v4.0.3.

#### Data alignment and read counting

We obtained single cell RNA-seq data from 429 samples containing 399 neurons and 30 negative controls. Negative controls contained sample collection solution (5% RNase inhibitor, 0.2% Triton-X RNase-free PBS) prepared on different days of experiment. Alignment was done using STAR (v.2.7.9a) on GRCm39 and Gencode vM27 with STAR parameters: --outFilterMultimapNmax 1 -- outSAMstrandField intronMotif --outFilterIntronMotifs RemoveNoncanonical. We retained 352 neurons with > 26000 unique aligned reads (median negative control: 26299.5) and > 5% fraction uniquely aligned reads (median negative control: 2.3%). Reads in exonic and intronic regions (Gencode vM27) were counted using the aligned bam files produced by STAR and summarizeOverlaps (GenomicAlignments v1.26.0) with singleEnd=TRUE, mode = “IntersectionNotEmpty”, ignore.strand = T, inter.feature = T parameters. TPM values were calculated separately for exonic and intronic regions using calculateTPM (package scater v1.18.6). Intronic and exonic TPMs for each gene were summed up, transformed as log2(TPM+1) and used for creating a Seurat (v4.0.4) object with min.cells = 5, min.features = 500 parameters.

#### Normalized Marker Sum (NMS) analysis

Using the reference dataset we extracted read counts from dorsal midbrain (Tissue = MBd) cells, removed cells with duplicated CellIDs and created a Seurat object with min.cells = 5, min.features = 500. Expression values were normalized (NormalizeData with normalization.method = “LogNormalize” and scale.factor = 10000 parameters) and scaled (ScaleData from Seurat package) using all genes. We identified top 200 marker genes for cell types from Taxonomy Rank 3 with more than 100 informative cells (Oligodendrocytes, Di- and mesencephalon neurons, Astroependymal cells, Vascular cells, Immune cells). Such analysis identified 986 unique genes (minimum of 188 genes per group). Next, we determined the highest expressing cell type for each marker gene to assign each gene to one cell type. We used the top 188 highest expressed genes for each group as marker genes to calculate the mean expression of cell-type marker genes in the corresponding cell type of the reference (mean expression value of all cells in the cell type). For each MADM-CloneSeq cell, the mean marker gene expression for each reference cell type was determined (5 expression values for each MADM-CloneSeq cell). Finally we determined the NMS score for each cell type by dividing the mean marker gene expression in each MADM-CloneSeq cell by the mean expression of cell type in the reference. Note that the absolute gene expression levels of the reference and the MADM-CloneSeq cells were of a different scale due to the difference in the calculation of gene expression (TPM vs UMI counts). More specifically the MADM-CloneSeq TPM gene expression values were ∼8x higher than the corresponding UMI counts in the reference. Since the NMS score is a direct comparison of reference / MADM-CloneSeq cells, we changed the neuron NMS cutoff from 0.4 (Lee et al., 2021) to 3. We thus only retained MADM-CloneSeq cells with a neuron NMS score > 3 and the highest non-neuronal NMS score < 4 (312 neurons). Finally, we corrected for possible batch effects in the MADM-CloneSeq data using SCTransform with vars.to.regress = c(“Plate”, “STAR.perc”) and residual.features = [all genes in expression matrix] parameters.

#### Data integration and cell type assignment

##### Reference cells

We extracted 16 cell types (MEGLU1, MEGLU2, MEGLU3, MEGLU4, MEGLU5, MEGLU6, MEINH2, MEINH3, MEINH5, MEINH6, MEINH7, MEINH8, MEINH9, MEINH10, MEINH11, MEINH12) that were located in our region of interest based on the predicted positional location (http://mousebrain.org/celltypes/). We extracted 3696 cells from these cell types (using column attribute ClusterName in the reference loom file) and created a Seurat object with min.cells = 5, min.features = 500 parameters. To prepare a reference UMAP we used SCTransform with standard parameters, RunPCA (assay = “SCT” parameter) and RunUMAP (dims = “1:20” parameters). For data integrations, reference data was processed using SCTransform with all genes of the count matrix as residual.features. Then we identified marker genes for each cell type using FindAllMarkers with logfc.threshold = 0.2 parameter. Top 100 marker genes for each cell type were determined and intersected with informative genes in the scale.data slot of the SCT assay of both reference and MADM-CloneSeq cells. Such analysis identified 996 genes that were used as features for data integration. For data integration we used the scale.data slot of the SCT assay by modifying the RunFastMNN function of the seurat-wrappers package. Note that RunFastMNN is a wrapper for the fastMNN function of the package batchelor (v1.6.3). For cell type assignment we used the expression matrix reconstructed from the low-rank approximation from fastMNN integration (assay = “mnnreconstructed”). We calculated the median expression of the same 996 genes identified above to calculate the mean expression across all reference cells in each cell type (centroid). Then we determined the Pearson correlation and associated test statistic (function cor.test with alternative = “two.sided”, method = “pearson” parameters) of each MADM-CloneSeq cell to each centroid. Each MADM-CloneSeq cell was assigned the cell type to which it had the highest correlation (nearest centroid). To assess the robustness of the mapping we used bootstrapping over genes. We selected 100 bootstrap samples of 996 genes with replacement, performed the nearest centroid classification and calculated a score by determining the fraction of the cell type assignment from bootstrapping analyses being identical to the cell type assignment of the real data (bootstrap score). For final analysis we used MADM-CloneSeq cells with highest correlation > 0.1, a bootstrap score > 0.5 and that belong to a clone with >30% collected cells (253 informative neurons). See Supplementary Table 1 for details on sample metadata.

##### UMAP visualization

To visualize MADM-CloneSeq cells on reference UMAP space we used a previously introduced pipeline (Berg et al., 2021) that uses Seurat as well as an R implementation of the UMAP library (https://github.com/tkonopka/umap) with modifications. Specifically, we used dimensionality reduction from fastMNN (first 25 dimensions) as well as the reference UMAP coordinates calculated above as an input into the UMAP pipeline.

##### Excitatory/inhibitory neuron marker analysis

Excitatory neurons were identified based on their respective cell types with excitatory neuron IDs starting with MEGLU and inhibitory neurons starting with MEINH. Note that for the sake of clarity we changed the cell type IDs for the MADM-CloneSeq cells in the final figures. For an exchange table see Fig. S1e. To identify differentially expressed genes (DEGs) between excitatory/inhibitory neurons in reference as well as MADM-CloneSeq data we used FindMarkers with parameter: logfc.threshold = 0.2. Genes were filtered for DEGs present in both analyses (reference and MADM-CloneSeq) and with an adjusted *p*-value < 0.01 in the MADM-CloneSeq data. A DE score was determined as the log10 of the adjusted *p*-value and corrected to be positive for genes with log2 fold-changes > 0. DE scores were cut at 5/-5 for better visualization, and plotted as a heatmap.

##### Cell-type marker analysis

We identified cell-type specific marker genes in both the reference and the MADM-CloneSeq data using FindAllMarkers with parameter: logfc.threshold = 0.2. We then determined the top 20 marker genes from the reference (ordered by average log2 fold-change) and intersected this gene list with the marker genes from the MADM-CloneSeq data. For heatmap visualization we prepared a differential expression score (DE score) by log2 transforming the raw *p*-value and correcting this value to be positive for genes with a log2 fold-change > 0. This value was cut at 30 for MADM-CloneSeq or at 200 for reference data. In Fig. S6i we used the whole gene list and for Fig. S6j we arbitrarily selected specific genes for display.

##### Relative abundance of cell types

To test for differences in the relative distribution of cell types between *Fzd10* and *Sox2* clones we plotted the 95% Clopper-Pearson confidence intervals (function clopper.pearson.ci from the GenBinomApps package v1.1). Largely overlapping intervals indicated non-significant differences, which we confirmed using chi-square test with correction for multiple testing (all adjusted *p*-values = 1, function chisq.test with parameters: correct = F, rescale.p = F, simulate.p.value = T).

##### Layer-specific cell-type distribution

To identify layer specific cell-type distribution we determined the number of neurons of each cell type in each layer (Subregion column of Supplementary Table 1). To test difference to a random distribution we permuted the Subregion column 1000 times and calculated a z-score. The highest z-score for each cell type was used to calculate a p-value using pnorm function.

To determine a random distribution of cell types in clones we considered hypothetical clones with 2, 3, 4, 5, 6, 7, 9, 10 and 12 cells. Then we randomly assigned one of the 16 cell types for each cell per clone. For each of these hypothetical clones we determined the number of unique cell types assigned to it. Randomization was done 1000x to calculated mean and standard deviation of unique cell types over all randomization. This data was used to draw the area of random distribution in Fig. 3l as well as to calculate a z-score with associated *p*-value (using pnorm function). Note that after correction for multiple testing (p.adjust function with method = “bonferroni”) all adjusted *p*-values = 1.

For preparation of the cell type linkage heatmap we determined a binary matrix indicating whether the respective cell type was assigned to one or more cells of the respective clone (black color) with rows representing cell types and columns representing clones. Heatmap and columns clustering was performed using pheatmap (v1.0.12) with parameters: clustering_distance_cols = “binary”, clustering_method = “ward.D”. Note that we restricted analysis for Fig. S7m - S7n to 13 cell types with a significant association with a single layer. Fig. S7j and S7n: AU (Approximately Unbiased) was determined using the same matrix as above using the pvclust package (v2.2.0, parameters: method.dist = “binary”, method.hclust=“ward.D”, nboot = 10000). Note that an AU >95% is typically used to determine clusters that are strongly supported by data (Suzuki and Shimodaira, 2006).

### *Pten* scRNA-sequencing data analysis

#### Reference data

For *Pten* control and *Pten* KO cell isolation, a *Nestin*-Cre driver (Petersen et al., 2002) with broad expression (i.e. not specific to midbrain) was used. Therefore we allowed for the possibility that the dissected midbrain contained “contaminating” cells from surrounding tissues, most likely hindbrain. To identify and remove non midbrain cells we established an embryonic reference following similar principles as described for Fig. 1 but with different filters. We focused on e18 cells, since this age is closest to P0 data analyzed here. Specifically cells with Tissue tag Midbrain and Hindbrain as well as Age tag e18 were extracted. No other filters were used to allow for a broad annotation of cell types based on this embryonic reference. Workflow: CreateSeuratObject with min.cells = 5, min.features = 500 parameters. NormalizeData with normalization.method = “LogNormalize”, scale.factor = 10000 parameters. FindVariableFeatures with selection.method = “vst”, nfeatures = 2000 parameters.

#### Pten control and Pten KO cells

Data from sorted Control-MADM and *Pten*-MADM was processed using cellranger-7.0.0 with transcriptome version: mm10-2020-A, including intronic information. We created a Seurat object from the filtered_feature_bc_matrix.h5 file using CreateSeuratObject with min.cells = 3, min.features = 200 parameters and retained only high quality cells: nFeature_RNA > 200 & nFeature_RNA < 6000 & percent.mt < 5. Data normalization and variable feature detection was done for each of the 2 datasets as for the embryonic reference. Transfer anchors were determined between the embryonic e18 reference and the Control-MADM/*Pten*-MADM data using FindTransferAnchors with dims = 1:30 parameter. These anchors were used to transfer the Class and Tissue labels from embryonic e18 reference to Control-MADM/Pten-MADM cells using TransferData with dims = 1:30 parameter. For further analyses Control-MADM and *Pten*-MADM data was integrated using: SelectIntegrationFeatures, FindIntegrationAnchors, IntegrateData. Filtering steps: Starting number of high quality cells: 4212/4713 (Control-MADM/*Pten*-MADM). Remove cells with predicted Tissue hindbrain: 3049/1982. Remove cells with predicted Class Immune, Fibroblast, Vascular, Subcommissural organ, Blood: 2985/1889.

#### Further analysis

After filtering the combined Control-MADM/*Pten*-MADM data was further processed: ScaleData, RunPCA(npcs = 40), RunUMAP(reduction = “pca”, dims = 1:35), FindNeighbors(reduction = “pca”, dims = 1:35), FindClusters(resolution = 0.5). Fig. 4o: UMAP with Class attribute as colors. Since Radial glia and Ependymal cells are likely producing glia cells at this developmental stage, we renamed these cell types as glia progenitors. Neuroblast was renamed immature neurons. Fig. 4p: Cells with Class tag Neuron and Neuroblast were extracted and relative abundances calculated. Error bars were calculated as clopper.pearson.ci with alpha = 0.05, CI = “two.sided” parameters. Fig. 4q: Differential expression statistics were calculated using FindMarkers ident.1 = Pten-MADM, ident.2 = Pten-Control, test.use = “MAST”, logfc.threshold = 0.1 separately for neurons and neuroblasts/immature neurons. MAST was v1.22.0. DEGs were defined as adjusted *p*-value < 0.01 and avg_log2FC > 0 (up regulated) or avg_log2FC < 0 (down regulated). Fig. 4r: Gene Ontology enrichment analysis of DEGs was performed with enrichGO (clusterProfiler v4.4.4) and OrgDb = org.Mm.eg.db (v3.15.0), ont = “ALL”, readable = T parameters. The number of significantly enriched GO terms (adjusted *p*-value < 0.05) falling into 3 categories was determined, using the search pattern given in brackets: Interneuron (GABA|interneuron|inhibitory), Development (differentiation & neuron), Cell cycle (cell cycle). Fig. S8f – S8g: DEG scores of genes mapping to one or more of the GO terms in the indicated group were determined as the log10 adjusted *p*-value. Prefix of the score was corrected to reflect direction of fold-change. Fig. 4s: Cells with the Class tag Neuron were extracted and processed: ScaleData, RunPCA(npcs = 40), RunUMAP(reduction = “pca”, dims = 1:30), FindNeighbors(reduction = “pca”, dims = 1:30), FindClusters(resolution = 0.5). Actual UMAP with gene expression was prepared using FeaturePlot with features = c(“Gad2”, “Slc17a6”), blend = T, order=T, min.cutoff = “q10”, max.cutoff = “q95”, cols = c(“#6d90ca”, “#d8a428”) parameters. Corresponding legend is shown in Fig. S8j. Fig. S8h – S8i: Cells were classified by investigating expression of 4 marker genes in the Seurat clusters. The association of Seurat clusters with excitatory (exc), inhibitory (inh) or mixed (exc/inh) cell type is as follows: 1: exc, 2:exc/inh, 3: exc, 4-7:inh. Fig. 4t: Relative abundance of exc, inh and exc/inh cell types was calculated. Error bars were calculated as clopper.pearson.ci with alpha = 0.05, CI = “two.sided” parameters. Fig. 4v: The combined embryonic pool/*Fzd10*-lineage data from Fig. 1 was used as a reference to transfer the labels of the Seurat clusters as shown in Fig. S1c and S2q to Control-MADM and *Pten*-MADM data: FindTransferAnchors (dims =1:30), TransferData (dims=1:30). Note that we focused this analysis on clusters containing mainly neurons (excluding clusters 0, 6, 11). Based on this cluster annotation we calculated relative abundances separately for Control-MADM and *Pten*-MADM data and determined significance of difference between Control-MADM and *Pten*-MADM. Only clusters with significant differences are shown. All significances between relative abundances were calculated using chisq.test with correct = T, rescale.p = F, simulate.p.value = F and corrected for multiple testing using p.adjust with method = “BH” parameter. Stars: *** < 0.001, ** <0.01, * < 0.1.

#### Velocity analysis

For velocity analysis from Seurat object we used the embryonic data from this study (Fig. S2i – S2o) and followed https://github.com/basilkhuder/Seurat-to-RNA-Velocity using velocyto v0.17.17 and scvelo v0.2.4.

### Statistical analysis

Data for clonal analysis, histological studies and MADM-CloneSeq documentations were stored and processed using Microsoft Excel (Microsoft). Statistical analyses were performed using GraphPad Prism software v8 (USA). All data were expressed as mean or median ± standard error of the mean (SEM) or upper and lower limits where n represents the number of clones, unless otherwise stated. To test for normal distribution, Shapiro-Wik normality test was used. To test for statistical significance between two groups, two-sided Mann-Whitney test was used. To test for statistical significance between more than two groups, one-way ANOVA was performed with Dunn’s multiple comparisons test between groups. Two-way ANOVA was used to compare the interactions between two sources of variations and to compare histogram distributions with Tukey’s multiple comparisons test used to test between groups. When comparing changes in relative abundance across different conditions, Fischer’s exact test was used. In some cases, one phase decay exponential curve fit was performed to determine R^2^ goodness of fit values. All statistical tests and p-values for scRNA-seq analysis are detailed in methods and/or corresponding figure legends.

## Acknowledgements

We thank Liqun Luo for his continued support, for providing essential resources for generating *Fzd10*-CreER mice which were generated in his laboratory, and for comments on the manuscript; W. Zhong for providing *Nestin-*Cre transgenic mouse line for this study; A. Heger for mouse colony management; R. Beattie and T. Asenov for designing and producing components of acute slice recovery chamber for MADM-CloneSeq experiments; and K. Leopold, J. Rodarte and N. Amberg for initial experiments, technical support and/or assistance. This study was supported by the Scientific Service Units (SSU) of IST Austria through resources provided by the Imaging & Optics Facility (IOF), Laboratory Support Facility (LSF), Miba Machine Shop, and Pre-clinical Facility (PCF). G.C. received funding from European Commission (IST plus postdoctoral fellowship). This work was supported by ISTA institutional funds; the Austrian Science Fund Special Research Programmes (FWF SFB F78 Neuro Stem Modulation) to S.H.

## Author contributions

G.C., F.M.P. and S.H. conceived the research. G.C., F.M.P. and S.H. designed all experiments and interpreted the data. G.C., F.M.P., P.K., T.K., C.S., M.S., N.G., and A.E.I., performed all the experiments. S.H., C.B. and R.S. provided reagents and/or resources. S.H. supervised the project. G.C., F.M.P. and S.H. wrote the manuscript. All authors edited and proofread the manuscript.

## Data and materials availability

All data generated and analysed in this study are included in the paper and/or supplementary materials. Raw sequencing data will be submitted to Gene Expression Omnibus (GEO).

## Code availability

All scripts that were used to prepare data and figures for this manuscript will be made available via GitHub. Files for 3D printing of acute slice recovery chamber (3D model 361319) are available under the following link: https://www.printables.com.

## SUPPLEMENTARY FIGURE LEGENDS

**Figure S1.**
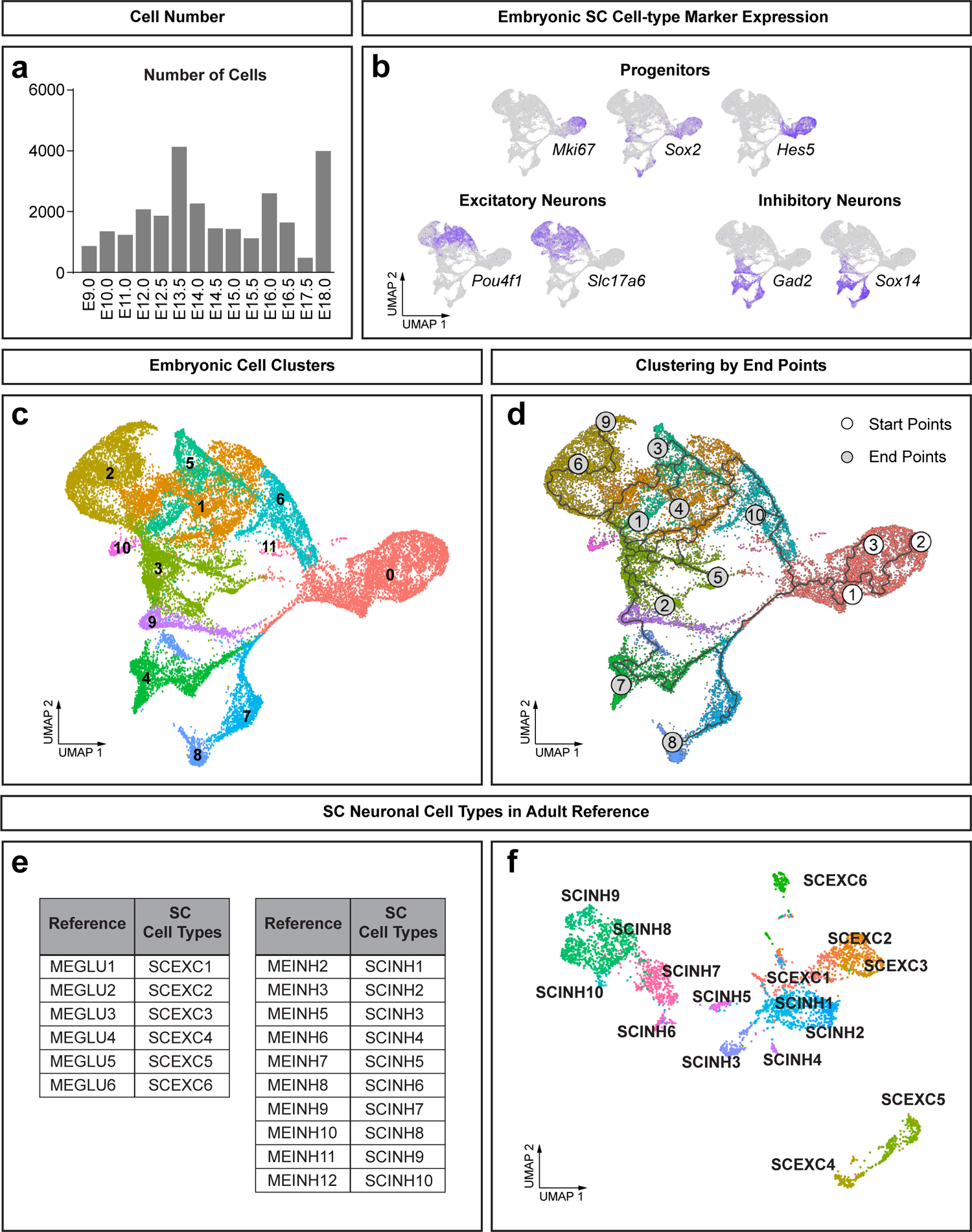
scRNA-seq data analysis of the embryonic and adult SC. **(a)** Summary of the number of cells extracted from La Manno embryonic dataset (La Manno et al., 2021). **(b)** Expression of cell-type markers in progenitors, excitatory and inhibitory neurons depicted on UMAPs of embryonic cell types. **(c)** UMAP showing 11 embryonic cell clusters as defined by unsupervised cluster analysis. **(d)** Overlay of embryonic development trajectories over 11 distinct cell clusters in the embryonic SC. **(e-f)** 16 adult SC neuronal cell types extracted from Zeisel adult dataset (Zeisel et al., 2018) were denominated accordingly (e) and formed distinct clusters on UMAP (f).

**Figure S2.**
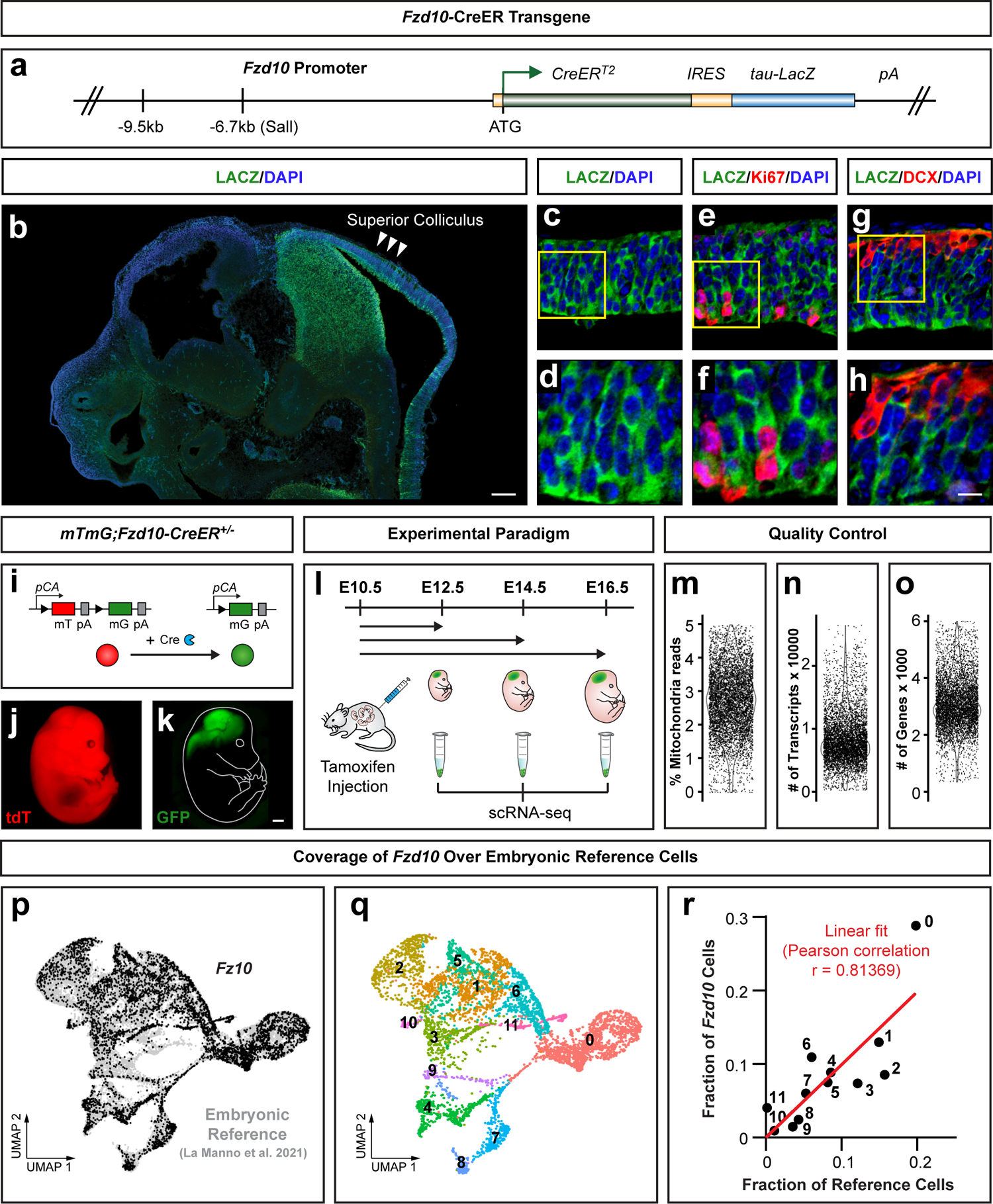
Generation and validation of *Fzd10-*CreER driver targeting SC progenitors. **(a)** Illustration of the *Fzd10*-*CreER* transgene. **(b)** Representative image indicating *Fzd10-CreER-TLZ* transgene expression at E11.5 in an overview. LACZ expression is evident in the midbrain, prospective superior colliculus is indicated. **(c-h)** Representative images illustrating *Fzd10-CreER-TLZ* transgene expression, as indicated by LACZ (green) signal (c and d) in proliferating Ki67^+^ progenitors (e and f) but less in nascent DCX^+^ postmitotic cells (g and h) in the dorsal midbrain at E10.5. Magnified views (d, f, and h) of corresponding yellow boxes in (c, e, and g) are indicated. **(i)** Schematic diagram illustrating *Cre*-mediated recombination that results in fluorescent color switch of the *mTmG* reporter. **(j and k)** Representative images of *mTmG;Fzd10-CreER^+/-^;* embryo injected with tamoxifen at E10.5 and imaged at E14.5. Cells that did not express *Fzd10-CreER^+/-^*were tdT^+^ (red) (j) whereas cells expressing *Fzd10-CreER^+/-^*were GFP^+^ (i.e. recombined *mTmG* reporter) (k). **(i)** Sample collection of cells within *Fzd10^+^* lineage, labelled at E10.5 by TM administration in *mTmG;Fzd10-CreER^+/-^*, and isolated from the embryonic dorsal midbrain at E12.5, E14.5 and E16.5. **(m-o)** Dot and violin plots showing quality control statistics of the percentage of mitochondria reads (m); the number of transcripts (n); and the number of genes (o) in all *Fzd10*-lineage cells used for the analysis. **(p and q)** UMAPs showing *Fzd10*-lineage cells in overlay with La Manno dataset (p) or defined as 11 cell clusters according to Fig. S1c (q) **(r)** Quantitative assessment of the similarity between *Fzd10*-lineage and La Manno dataset, based on the correlation of relative distribution of each cell cluster (red line = linear fit with Pearson correlation). Scale bars = 100μm (b); 20μm (c, e, and g); 10μm (d, f, and h); 1mm (j and k).

**Figure S3.**
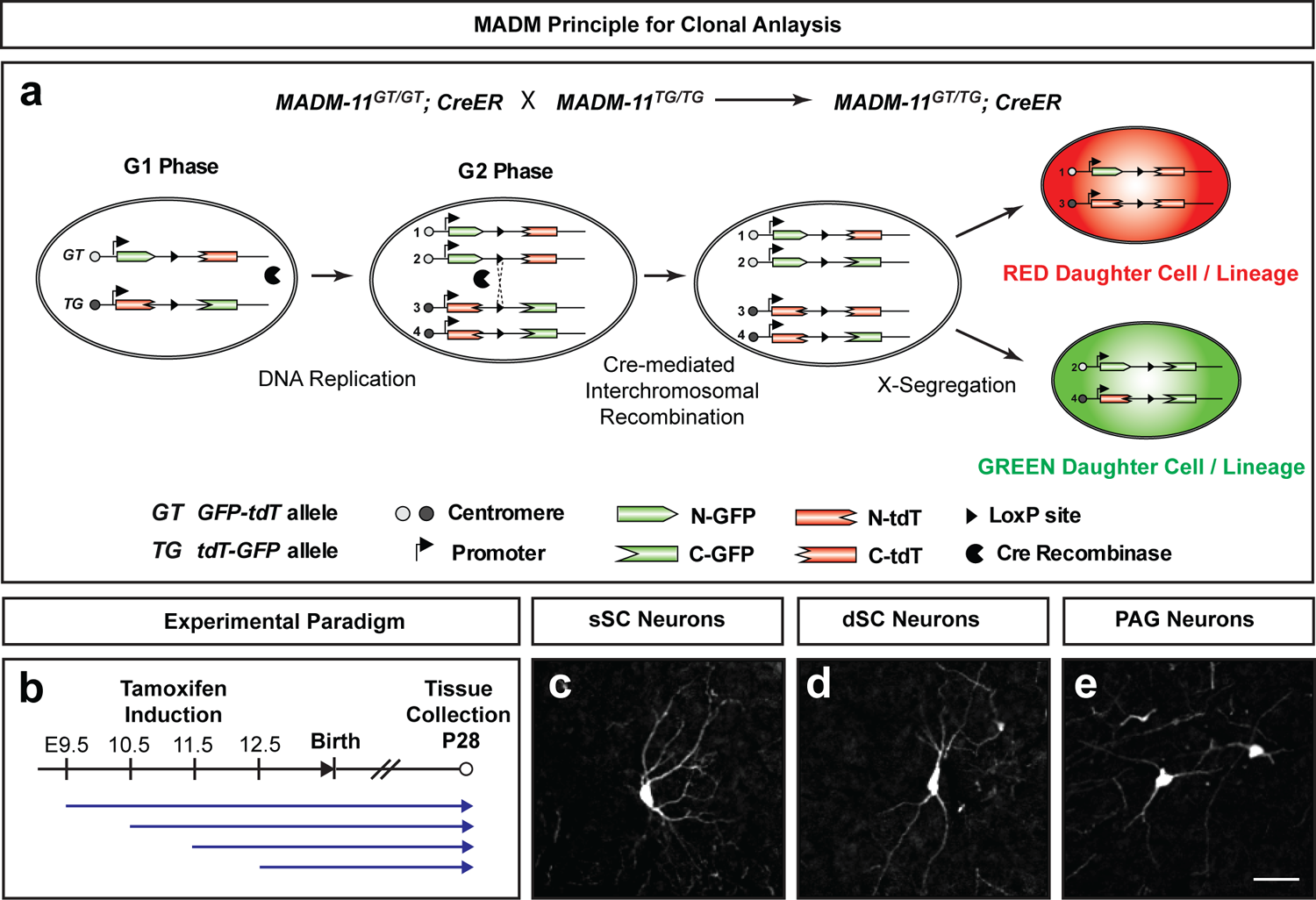
MADM clonal analysis in the SC. **(a)** Schematic overview of the MADM principle resulting in differential labelling of stem cell derived daughter cells and their respective lineages marked in red (tdT) and green (GFP). **(b)** Experimental MADM clonal labelling paradigm. TM-based MADM clone induction at E9.5, E10.5, E11.5 or E12.5 developmental time points with tissue collection and analysis at P28. **(c-g)** Representative images of MADM-labelled neurons in the sSC (c), dSC (d), and PAG (e). Scale bar = 20μm (c-e).

**Figure S4.**
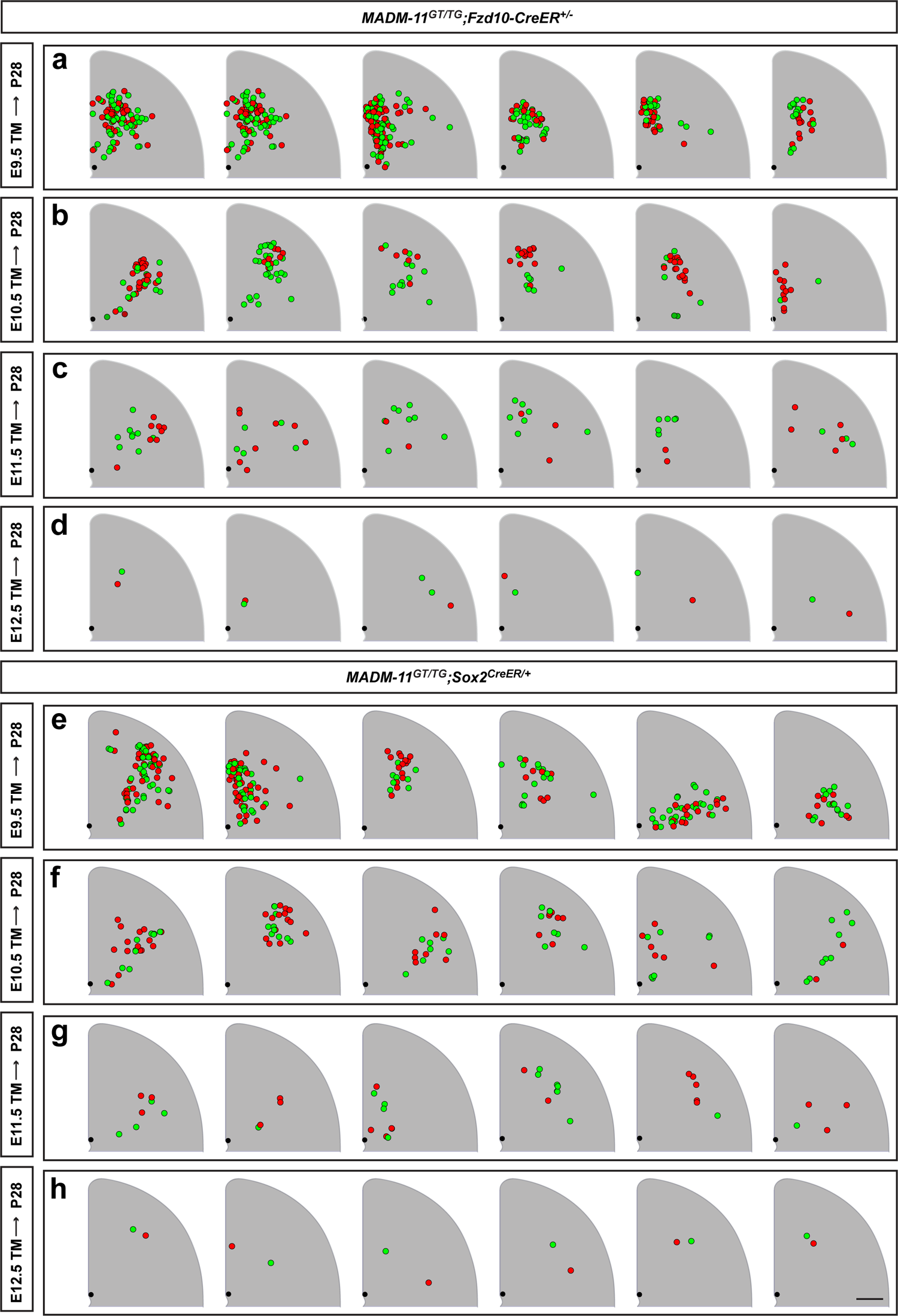
Reconstructions of MADM clones in SC. **(a-h)** Reconstructions of SC MADM clones in *MADM-11^GT/TG^;Fzd10-CreER^+/-^*(a-d) and *MADM-11^GT/TG^;Sox2^CreER/+^* (e-h) with TM-mediated clone induction at E9.5 (a and e), E10.5 (b and f), E11.5 (c and g) and E12.5 (d and h) and analysis at P28. Each clone is illustrated as a maximum z-projected cluster of neurons shown as red and green dots. Precise locations of each cell relative to a reference point at the aqueduct (black dot) are plotted. Grey areas mark SC hemispheres. For illustration purposes, all clones found on the left hemispheres were flipped. Scale bar = 500μm.

**Figure S5.**
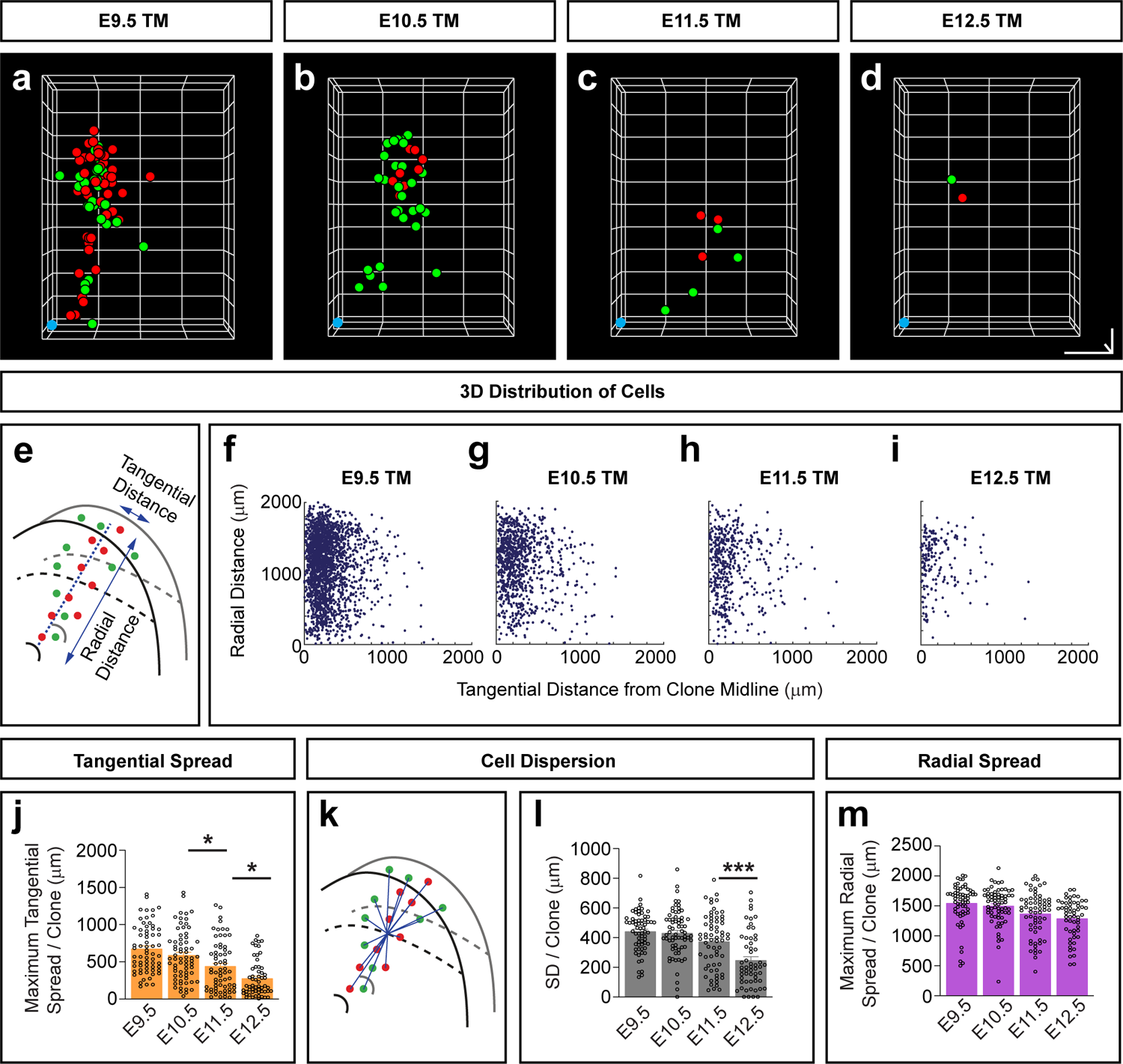
3D reconstruction of MADM clone architecture in SC. **(a-d)** Representative 3D illustrations of neuronal distribution in MADM clones in SC for TM induction time points at E9.5 (a), E10.5 (b), E11.5 (c) and E12.5 (d) with analysis at P28. Each dot represents the location of a single neuron in either red or green. Blue dot marks the location of the aqueduct (AQ) in the middle of the clone as a reference point. **(e)** Schematic illustration indicating the determination of radial and tangential distance of each cell in 3D from a geometric midline originating from the reference point (dotted line). **(f-i)** Plots of radial versus tangential distance of each cell depicted for E9.5 (f), E10.5 (g), E11.5 (h), and E12.5 (i) TM induction time points (n=2110 cells in f, n=925 cells in g, n=459 cells in h, n=179 cells in i). **(j)** Quantification of the maximum tangential spread in each clone across induction time points. **(k and l)** Illustration (k) and quantification (l) of cell dispersion in each clone determined by the standard deviation of the distance of each cell from the centroid of the clone. All bars indicate mean ± SEM. E9.5 (n=64), E10.5 (n=68), E11.5 (n=65), and E12.5 (n=57). **(m)** Quantification of the maximum radial spread in each clone across induction time points. One-way ANOVA with Dunn’s post-hoc test for comparisons between consecutive time points: *p*=0.3991, 0.0265, 0.0158 in j; *p*>0.999, *p*=0.257, 0.0009 in l; *p*=0.6195, 0.0835, 0.4076 in m. \**p*<0.05; \*\*\**p*<0.001. Scale bars = 500 x 200 x 1000 μm in x-, y- and z-dimensions (a-d).

**Figure S6.**
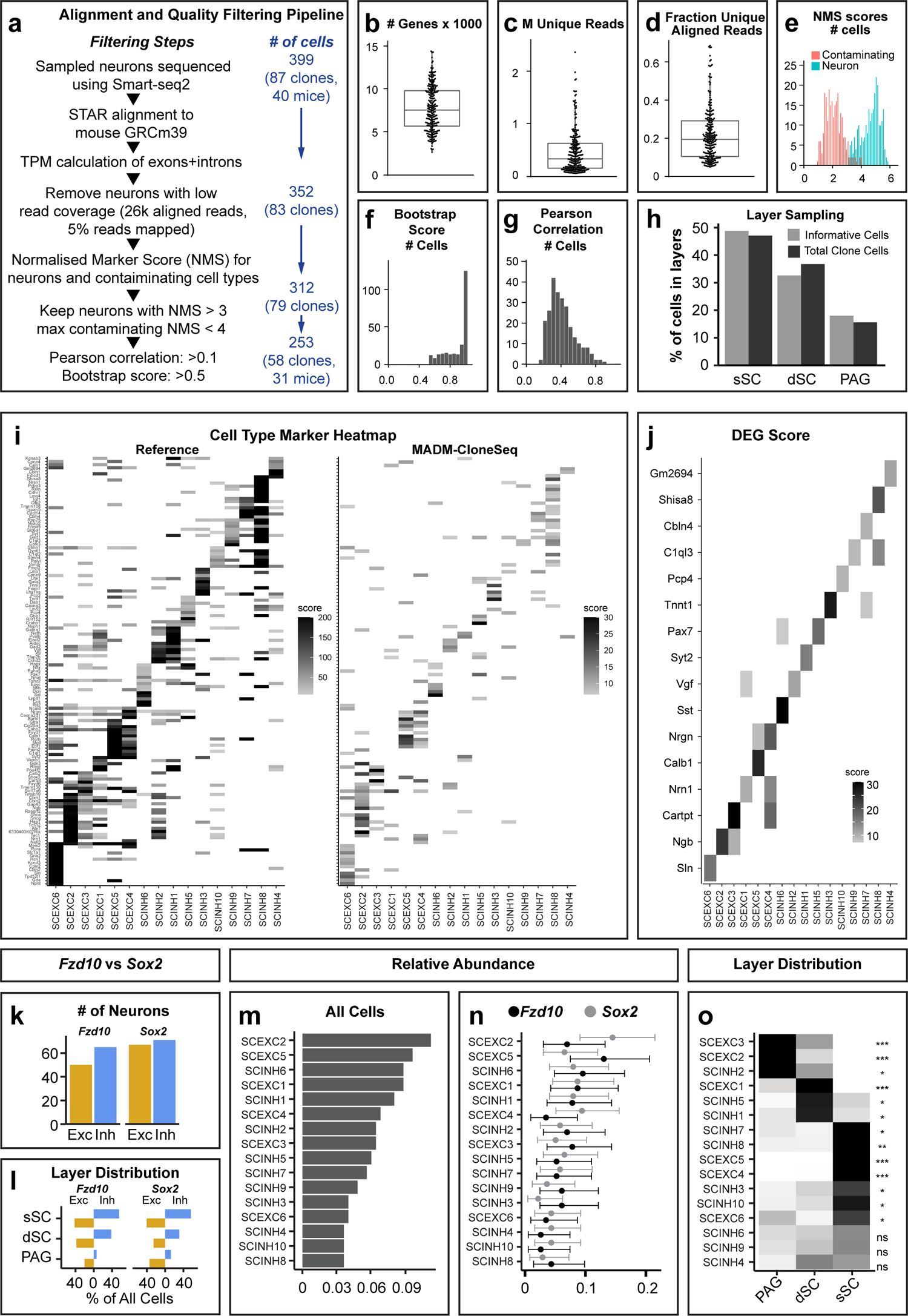
MADM-CloneSeq data analysis pipeline and quality control. **(a)** Workflow of RNA-sequencing data analysis of MADM-CloneSeq samples. **(b-d)** Quantification of genes (b), unique reads (c) and fraction unique aligned reads (d) detected per cell (box indicates median, 1^st^ and 3^rd^ quartile). **(e-g)** Statistics indicating the distribution of quality features of 253 high quality cells: NMS (e), bootstrap scores (f) and Pearson correlation (g). **(h)** Relative SC layer distribution of informative MADM-CloneSeq cells (grey bar) and total clone cells (black bar). No significant difference could be detected. **(i and j)** Heatmaps illustrating DEG scores of a list of marker genes (i) for reference (left) and corresponding MADM-CloneSeq (right) cell types; and a selected list of cell-type specific marker genes (j). **(k and l)** Quantification of the number of informative excitatory and inhibitory neurons (k) and their layer distribution (l), plotted separately for either *Fzd10-* or *Sox2*-CreER driver. **(m and n)** Quantification of the relative abundance (fraction ± 95% Clopper-Pearson confidence intervals) of different cell types plotted for both (m) and individual (n) *Fzd10-* or *Sox2*-CreER driver, respectively; for n=115 cells from *Fzd10* clones and n=138 cells from *Sox2* clones. No significant difference in relative distribution for any cell type was detected after multiple test correction (*padj*=1). **(o)** Heatmap indicating the layer-specific relative abundance of each cell type (z-score and associated *p*-values of lowest z-score per cell-type). Respective *p*-values from top to bottom: *p*=0.00000104, *p*=1.7×10^-12^, *p*=0.021, *p*=0.0000247, *p*=0.0424, *p*=0.0499, *p*=0.0131, *p*=0.00645, *p*=0.0000615, *p*=0.000545, *p*=0.0865, *p*=0.0419, *p*=0.096, *p*=0.308, *p*=0.242, *p*=0.221. ns = not significant, \**p*<0.1, \*\**p*<0.01, \*\*\**p<*0.001.

**Figure S7.**
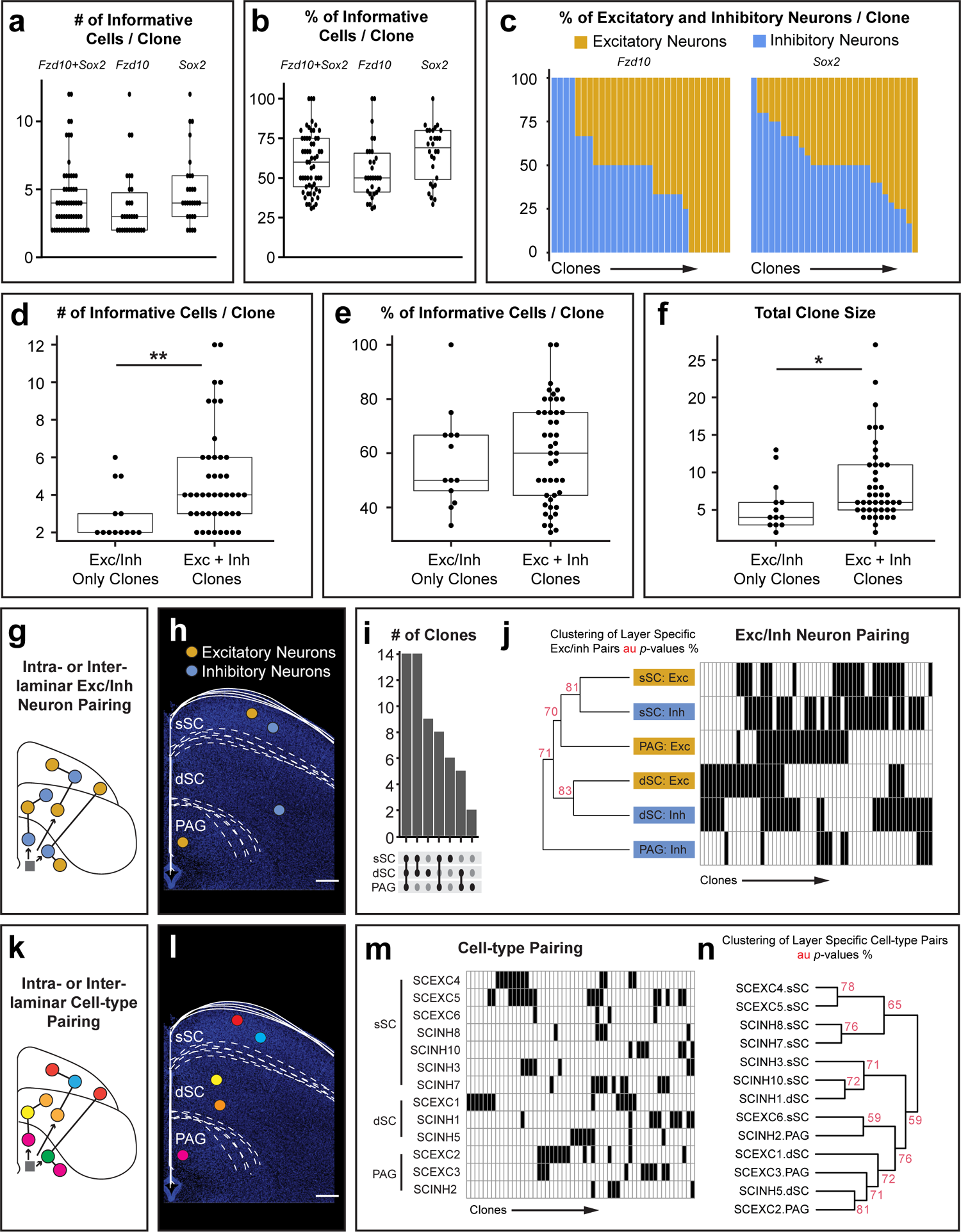
MADM-CloneSeq data provides evidence that SC cell type production exhibits no layer-restriction. **(a and b)** MADM-CloneSeq clonal sampling measured by the number (a) and the percentage (b) of informative cells per clone for both (left), *Fzd10* only-(middle) and *Sox2* only-(right) MADM clones (box shows median, 1^st^ and 3^rd^ quartile; n=30 *Fzd10* and n=28 *Sox2* clones). **(c)** Relative numbers of excitatory and inhibitory neurons in each MADM clone are indicated for *Fzd10* (left) and *Sox2* (right) clones. **(d-f)** Comparison of the number of informative cells per clone (d), clonal coverage (e), and total clone size (f) between clones containing only one principal neuron type (Exc/Inh only clones; n=13), and those containing both (Exc + Inh clones; n=45). Welch Two Sample t-test (two sided) for comparisons: *p*=0.002305 (d), *p*=0.7527 (e), and *p*=0.02759 (f). All boxes show median, 1^st^ and 3^rd^ quartile. **(g and h)** Schematic diagrams illustrating potential intra- or inter-laminar excitatory/inhibitory neuronal co-production with a hypothetical clone depicted in (h). **(i)** Number of clones containing cells in specific layer combinations used in subsequent analysis. **(j)** Presence of excitatory and inhibitory neurons in each layer shown as black boxes for each clone. Cluster analysis indicated no significant pattern of co-production of excitatory/inhibitory neuron types across layers (Approximately Unbiased *p*-value < 95%). **(k and l)** Schematic diagrams illustrating potential intra- or inter-laminar co-production of different SC neuron types with a hypothetical clone depicted in (l). **(m)** Presence of each SC neuron type in each layer was indicated as black boxes for individual clones. **(n)** Cluster analysis indicates no significant pattern of any neuronal type co-production (Approximately Unbiased *p*-value < 95%). \**p*<0.05, \*\**p*<0.01. Scale bar = 200μm (h and l).

**Figure S8.**
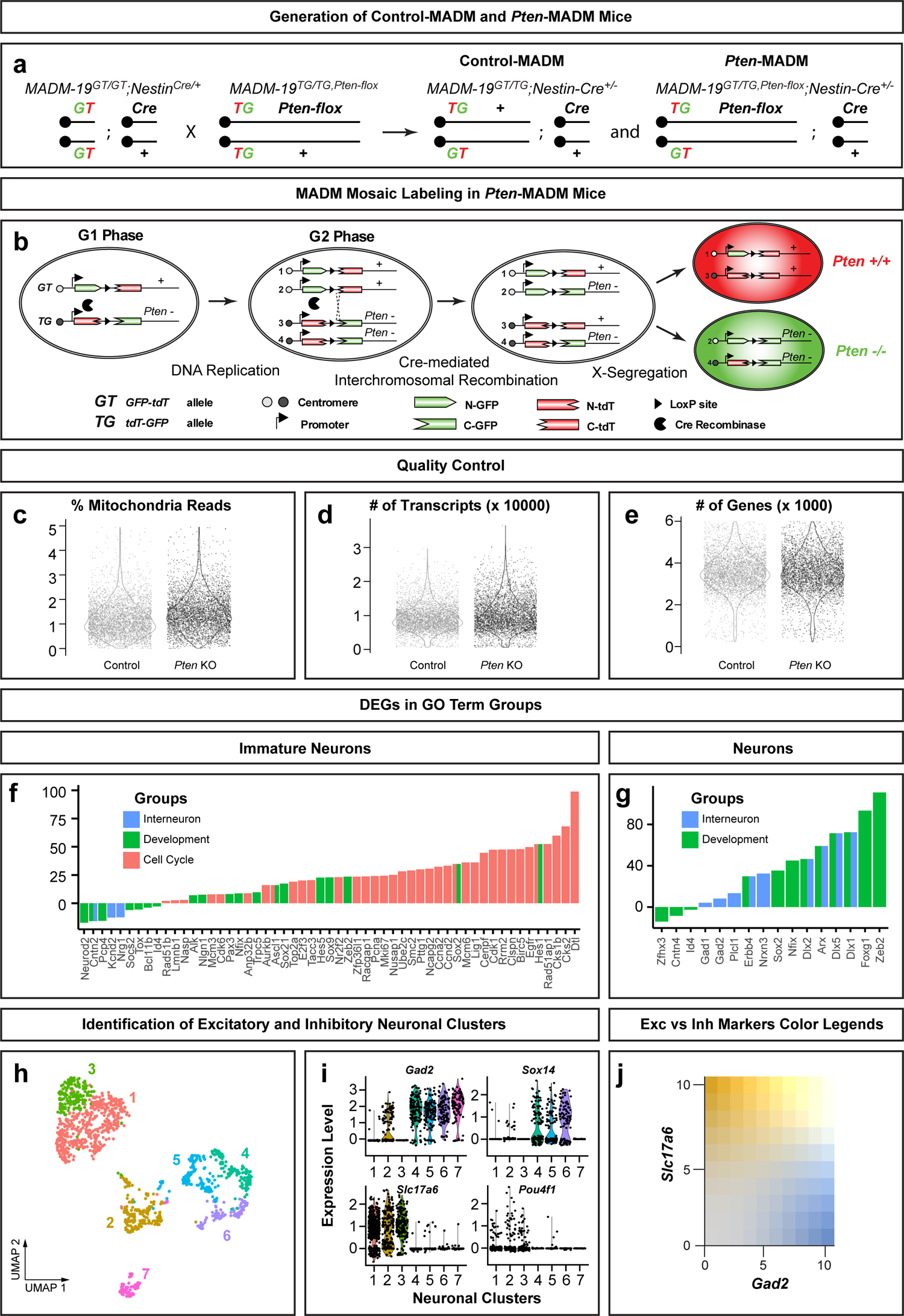
Generation and analysis of Control-MADM and *Pten*-MADM. **(a)** Breeding scheme for the generation of Control-MADM and *Pten*-MADM mice. **(b)** MADM principle resulting in mosaic labelling in *Pten*-MADM mice with red (tdT) *Pten^+/+^* and green (GFP) *Pten^-/-^* cells. **(c-e)** Dot and violin plots indicate quality control statistics of the percentage of mitochondria reads (c), the number of transcripts (d), and the number of genes (e) of each cell for Control and *Pten* KO. **(f and g)** Bar plots of DEG scores for genes within GO term groups for immature (f) and mature neurons (g). **(h and i)** Seven neuronal clusters depicted on UMAP (h) with their expression levels of inhibitory (*Gad2, Sox14*) and excitatory (*Slc17a6, Pou4f1*) subtype-specific markers (i). **(j)** Color legend corresponding to Figure 4s.

